# Selective representations of texture and motion in mouse higher visual areas

**DOI:** 10.1101/2021.12.05.471337

**Authors:** Yiyi Yu, Jeffrey N. Stirman, Christopher R. Dorsett, Spencer L. Smith

## Abstract

Mice have a constellation of higher visual areas, but their functional specializations are unclear. Here, we used a data-driven approach to examine neuronal representations of complex visual stimuli across mouse higher visual areas, measured using large field-of-view two-photon calcium imaging. Using specialized stimuli, we found higher fidelity representations of texture in area LM, compared to area AL. Complementarily, we found higher fidelity representations of motion in area AL, compared to area LM. We also observed this segregation of information in response to naturalistic videos. Finally, we explored how popular models of visual cortical neurons could produce the segregated representations of texture and motion we observed. These selective representations could aid in behaviors such as visually guided navigation.

## Introduction

Visual systems evolved to extract behaviorally relevant information from complex natural scenes. Visual stimuli contain information about texture, motion, objects, and other features of the environment around the animal. These components of visual stimuli have unequal relevance across behaviors. For example, optic flow and parallax motion information can help guide navigation behavior, but object recognition is often invariant to motion. The ventral stream of cortical areas in rodents function as object detection circuitry, as they do in primates. As expected, these areas exhibit neural representations (spatiotemporal patterns of neuronal activity, a.k.a. population codes) that are increasingly invariant in their responses with changes in the appearance of recognized objects^1–3^.

In mice, axons from neurons in primary visual cortex (**V1**) extend out to an array of higher visual areas (**HVAs**), seven of which share a border with V1, and all of which have characteristic interconnectivity with other brain regions. Mouse visual cortical areas exhibit a level of hierarchical structure, and form two subnetworks^4–9^. HVAs receive functionally distinctive afferents from V1 (ref. ^10^). At least nine HVAs exhibit retinotopic topology^8,11–13^ and neurons in HVAs have larger receptive fields than neurons in V1 (ref. ^8^). This organization and connectivity of mouse visual areas may have evolved to selectivity propagate specific visual information to other brain regions, but the functional specializations of HVAs require further elucidation.

Gratings are classic visual stimuli for characterizing responses in visual cortical areas^14–16^. In mice, HVAs exhibit biases in their preferred spatial and temporal frequencies of gratings, but overall, their frequency passbands largely overlap^10,17–19^. Similar studies using alternative visual stimuli have produced additional insights: spectral noise stimuli revealed further details of spatiotemporal preferences among HVAs^20^; plaid stimuli (two superimposed gratings with different angles) revealed pattern cells in LM and RL^21^; naturalistic texture stimuli were better discriminated from scrambled versions in LM than in V1^22^, and random dot kinematograms highlighted motion-coherent modulation in putative dorsal areas AL, PM, and AM^15,23^. One could hypothesize texture and motion to be key components of any visual stimuli. How are representations of texture and motion features in visual stimuli segregated among HVAs in mice? Representation of texture relies on the encoding of local features^24^. Experimental and theoretical studies suggested that HVAs may encode a combination of local features, such as multiple edges to detect curves and shapes^25–27^.

In the current study, we have examined the visual feature selectivity of multiple visual areas to three classes of visual stimuli: drifting textures, random dot kinematograms, and naturalistic videos. We have examined how the texture and motion components of a naturalistic video are represented, and found that high fidelity representations of these stimulus classes are segregated to different HVAs. We then explored how a range of popular Gabor filter-based models of visual cortical neurons can produce similar segregations of stimulus representations. The results from these experiments reveal new aspects of the tuning properties of mouse HVAs.

## Results

### Multi-area calcium imaging to distinguish tuning properties of HVAs

To survey the tuning properties of multiple visual cortical areas, we performed population calcium imaging of L2/3 neurons in V1 and four HVAs (lateromedial, **LM**; laterointermediate, **LI**; anterolateral, **AL**; posteromedial, **PM** or anteromedial, **AM**) of awake mice using a multiplexing, large field-of-view two-photon microscope with subcellular resolution developed in-house^28^, and transgenic mice expressing the genetically encoded calcium indicator GCaMP6s^29,30^. We located V1 and HVAs of each mouse using retinotopic maps obtained by intrinsic signal optical imaging^18,31^ (**Supplementary Fig. 1a**). Borders of HVAs were reliably delineated in most cases, with the exception being some experiments where the AM and PM boundary was not clearly defined (for those cases, neurons were pooled as AM/PM). We imaged neuronal activity in 2 – 4 cortical visual areas simultaneously (**Fig. 1a, b**). Calcium signals were used to infer probable spike trains for each neuron (*Methods*; **Supplementary Fig. 1b**). During visual stimulation, the average and maximal firing rates inferred were similar across cortical areas, and were typically around 0.5 spikes/s average, and ranged up to 15-30 spikes/s maximal (**Fig. 1c**).

**Figure 1.**
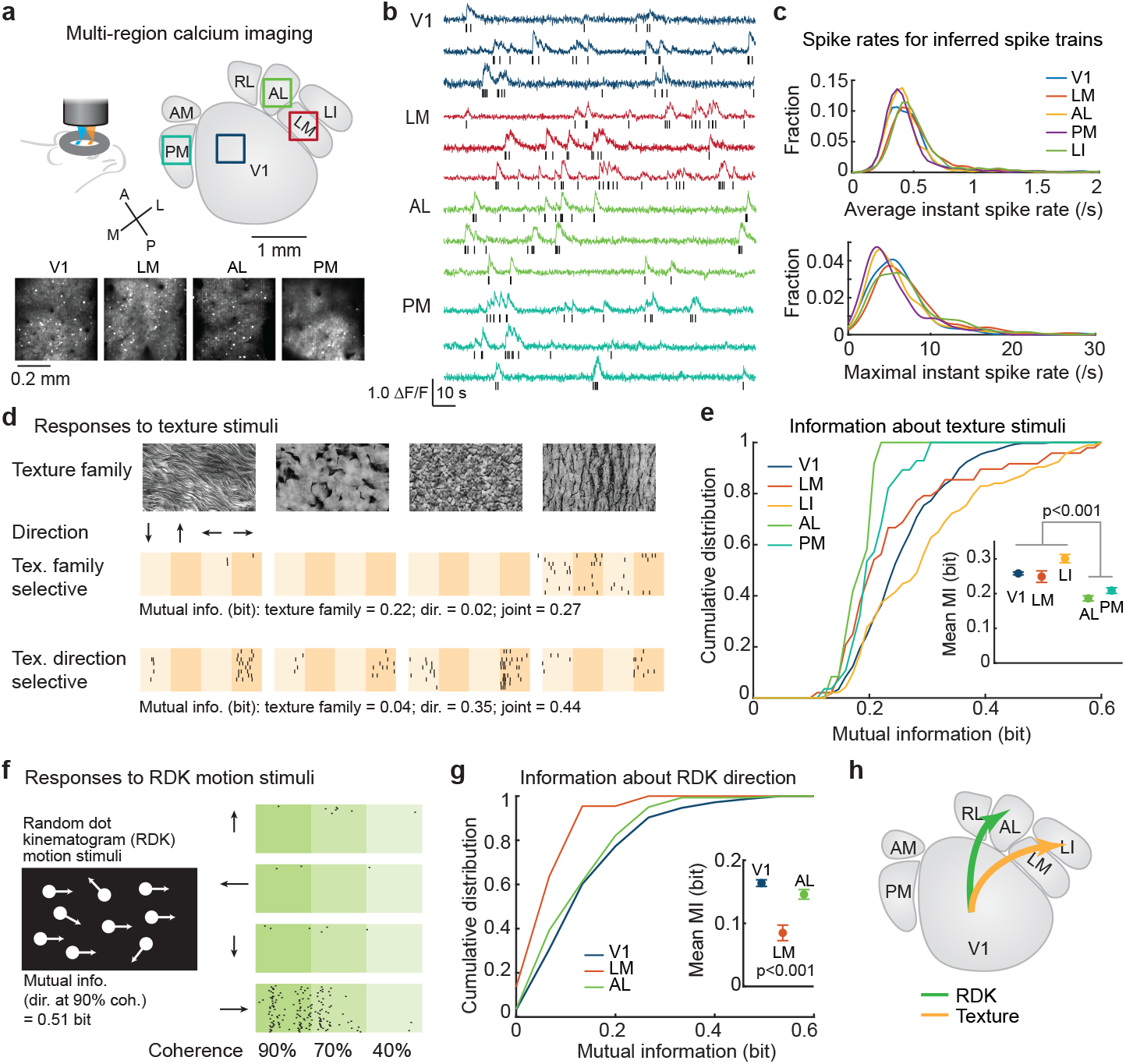
Segregated representations of textures and random dot kinematogram (RDK) motion in HVAs. (**a**) Neural activity was imaged in multiple HVAs simultaneously using large field-of-view, multiplexed two-photon calcium imaging. In an example experiment, layer 2/3 excitatory neurons were imaged in V1, LM, AL, and PM simultaneously. Squares indicate the imaged regions, and projections of raw image stacks are shown below. (**b**) Image stacks were analyzed to extract calcium dynamics from cell bodies, after neuropil subtraction. These traces were used to inferred spike activity, as shown in raster plots below each trace. (**c**) Statistics of inferred spiking were similar to those of prior reports, indicating accurate inference. The mean and maximal instantaneous firing rates of neurons in V1 and HVAs are similar (mean, 0.5 ± 0.5 spike/s; max, 7 ± 11 spike/s; p = 0.055; one-way ANOVA with Bonferroni correction). (**d**) Mice were shown texture stimuli, each of which was from one of four families, and which drifted in one of four directions. Spike raster plots from two example neurons (10 trials shown for each) show that one neuron is selective for texture family, and the other is more selective for texture direction. The amount of mutual information (**MI**, in bits) for the two stimulus parameters (texture family and panning direction) are written below each raster, along with the overall or joint (family and direction) MI. (**e**) V1, LM, and LI provide higher MI for texture stimuli than AL or PM (p = 5.8 x 10^-8^; one-way ANOVA, Bonferroni multiple comparison). Error bars in inset indicate SE. (**f**) Mice were shown random dot kinematogram (**RDK**) motion stimuli, which drifted in one of four directions with up to 90% coherence (fraction of dots moving in the same direction). A raster for an example neuron (30 trials) shows that it fires during rightward motion, with 0.51 bits of MI for motion direction at 90% coherence. (**g**) V1 and AL provide higher MI for the RDK motion direction than LM (p = 0.0006; one-way ANOVA, Bonferroni multiple comparison). (**h**) These results indicate a segregation of visual stimulus representations: texture stimuli to LM, and RDK motion stimuli to AL.

We characterized the neuronal responses to three types of visual stimuli: scrolling textures (hereafter “texture stimuli”), random dot kinematograms (**RDK**), and a naturalistic video mimicking home cage navigation (*For experiment details see* ***Table 1***). Neurons that fired on 60% of trials were considered “reliably responsive”, if not otherwise stated. In general, half of all recorded neurons responded to at least one visual stimulus reliably (texture: 55%; RDK: 54%; naturalistic video: 50%). For each stimulus type, we characterized the tuning properties of individual neurons using an encoder model (*Methods*). We also measured neuronal selectivity to texture family, motions direction, or joint selectivity using mutual information analysis. Higher bit values for a neuron-stimulus parameter pair means that the activity from that neuron provides more information about that stimulus parameter (or combination of stimulus parameters).

**Table 1.**
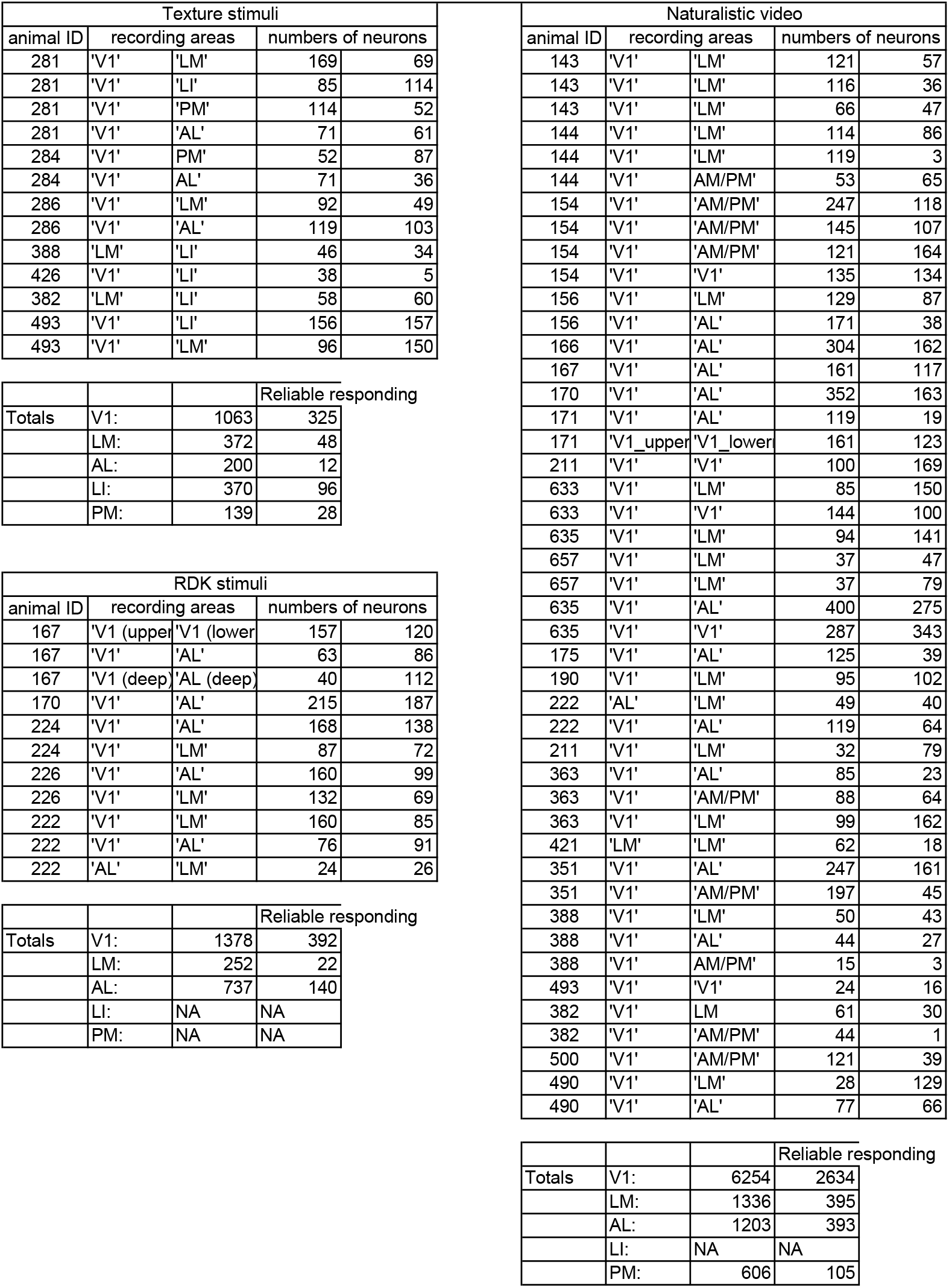
Summary of recording sessions.

### Information about texture and RDK were encoded in separate HVAs

We tested the selectivity of neurons in V1, LM, LI, AL and PM to texture stimuli using a set of naturalistic textures that drifting in one of the four cardinal directions (**Supplementary Fig. 2a**). We generated four families of texture images based on parametric models of naturalistic texture patterns (*Methods*). These stimuli allowed us to characterize the representation of both texture pattern information and drift direction information, and thus test the tolerance of a texture selective neuron to motion direction.

We observed reliable responses to drifting textures in V1, LM, LI and PM, while AL was barely responsive to these stimuli (**Supplementary Fig. 2b**). About 43% of reliably responsive neurons were modulated by the texture stimuli (i.e., texture-tuned neurons) (*Methods;* **Supplementary Fig. 2c-e**). Texture-tuned neurons exhibited various selectivity patterns, suggesting a variety of encoding properties (**Fig. 1d**). For example, about 13%-38% (varied across HVAs) of neurons were strictly selective to one texture family drifting in one direction (**Supplementary Fig. 2f**), and a different group of neurons (about 30%) were also selective to one texture family but responded to more than one motion direction of that texture family (**Supplementary Fig. 2f**). This latter group of tuned neurons could be called *tolerant* to motion direction, with the implication that it is selective for the other stimulus parameter (texture family, in this case). In general, we observed neurons tolerant to either texture family or motion direction in V1 and HVAs.

Using mutual information analysis, we then characterized the selectivity of individual neurons in HVAs. Overall, neurons in V1 and LI were more informative about the texture stimuli, followed by LM. By contrast, neurons in areas AL and PM were not informative about the texture stimuli (**Fig. 1e;** p = 5.8 x 10^-8^; one-way ANOVA). To examine the tolerance of texture encoding neurons to the translational direction, we computed the mutual information between neuronal responses and texture families (refer the statistical pattern of a texture image). LI was the most informative about texture family out of all tested visual areas, followed by V1 and LM (**Supplementary Fig. 3a;** p = 0.0006; one-way ANOVA). Meanwhile, V1 and LI also carried more information about the motion direction of the texture stimuli, compared to areas AL and PM (**Supplementary Fig. 3b;** p = 0.0006, one-way ANOVA). Examining the information encoding of individual neurons, we found an increasing fraction of neurons that jointly encoded texture family and texture drift direction along the putative ventral pathway (V1: 13%, LM: 17%, LI: 30%, **Supplementary Fig. 3c**; AL: 0%, PM: 0%), suggesting increasing joint coding along the putative visual hierarchy.

These results for texture encoding contrast with results for standard drifting gratings. For gratings, we found motion direction information to be encoded broadly, differing <10% among HVAs (**Supplementary Fig. 3d**), while texture motion information did not propagate to visual areas outside the putative ventral pathway, differing >250% among HVAs (**Supplementary Fig. 3a, b**). The drift speeds were similar (32 degrees/s for the textures and 40 degrees/s for the gratings), so it is unclear which spatial structural differences between these stimuli drove the differences in encoding. Thus, we next examined responses to a stimulus with less spatial structure and greater focus on motion.

We examined the encoding of random dot kinematograms (**RDK**), which are salient white dots on a dark background with 40-90% motion coherence (remaining dots move in random directions (**Fig. 1f; Supplementary Fig. 4a**). The RDK stimuli elicited responses in 40-80% of neurons in V1, LM and AL, and V1 and AL were more responsive to and generated more reliable representations of the RDK stimuli (**Supplementary Fig. 4b**). Among reliably responsive neurons (responding on at least 60% of trials), about 32-60% of neurons were modulated by the RDK stimuli (i.e., exhibited tuning) (V1: 59%, LM: 32%, AL: 43%; **Supplementary Fig. 4c, d**). RDK-tuned neurons exhibited selectivity to motion directions and were modulated by the motion coherence (**Supplementary Fig. 4e**). To characterize the direction selectivity, we computed the mutual information between neuronal responses and the motion direction at each coherence level (**Supplementary Fig. 4f**). We found that V1 and AL were more informative than LM about the motion direction of the RDK at all coherence levels (**Fig. 1g;** p = 0.0006, one-way ANOVA).

In summary, texture family selective neurons were found in V1, LM, LI and PM, while RDK direction selectivity neurons were more abundant in AL. Thus, information about drifting textures and RDK motion are segregated to distinct HVAs (**Fig. 1h**).

### Features of naturalistic videos were encoded in separate HVAs

To determine whether this segregation of texture and motion information among HVAs could be detected within a more complex stimulus, we characterized the cortical representation of a naturalistic video (**Fig. 2a**). The 64-second-long naturalistic video stimulus contained time-varying visual features such as contrast^32^, luminance, edge density^26^, difference of Gaussian (**DOG**) entropy^33^, and optic flow (**OF**) speed and direction^23^ (*Methods*; **Supplementary Fig. 5**). About 40% of neurons in the four areas responded to the naturalistic video reliably (trail-to-trial correlation > 0.08; **Supplementary Fig. 6a, b**). Our results thus far suggested that activity in AL would be modulated by motion information in the naturalistic video, and activity in LM would be modulated by texture information in the same video. We tested this hypothesis.

**Figure 2.**
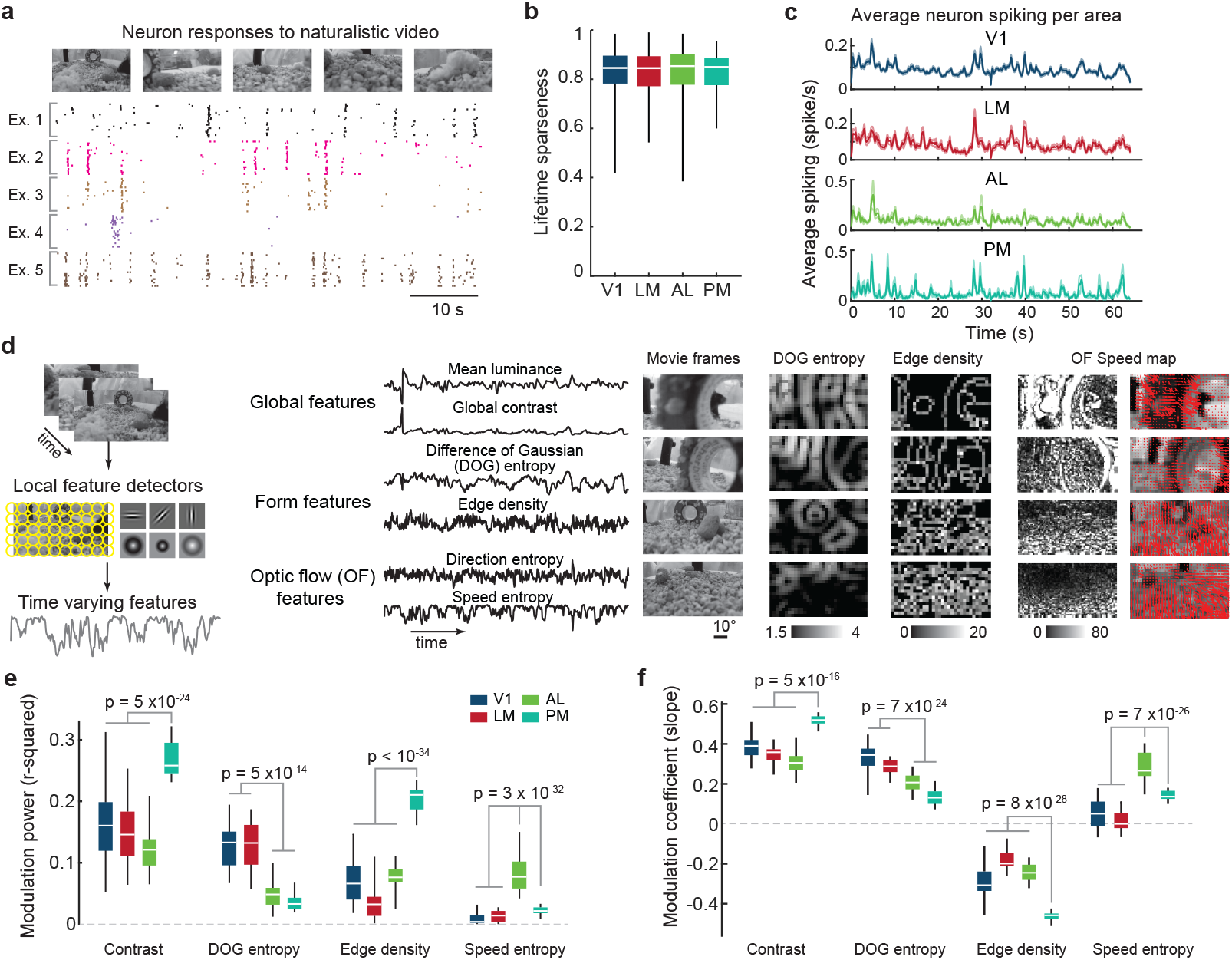
Segregated representations of spatial and motion features in naturalistic videos. (**a**) Five example neurons show reliable, yet diverse, spike responses during a naturalistic video stimulus. **(b)** Neurons in all four tested areas exhibited similarly high response sparseness to the naturalistic video (one-way ANOVA, p = 0.8). (**c**) Average spike responses varied across cortical areas (traces are averages, across reliable neurons). For example, neurons in area AL tended to show a spike in activity about 5 s into the video, whereas neurons in LM did not. Shaded area indicates SEM computed across multiple animals. (**d**) Form and motion components of the naturalistic video were extracted using a bank of linear filters with various sizes and locations (left). This provided time-varying signals correlated with global, form, and motion features, such as contrast, difference-of-Gaussian (**DOG**) entropy, and speed entropy (middle). To provide an intuitive feel for these features, example naturalistic video frames with the corresponding DOG entropy maps, edge density maps, and optical flow speed maps are shown (also see **Supplementary Fig. 5a, b**). (**e**) The time-varying features were weighted to best match the average neuronal activity for a cortical area (N = 200 with permutation). The linear weights are modulation coefficients and the goodness-of-linear fitting, or r-squared, is the modulation power. Areas V1 and LM was strongly modulated by DOG entropy, but AL and PM were not. Area LM was the only area modulated by speed entropy. Area PM was modulated by contrast and edge density. (**f**) The modulation coefficients were typically positive, but were negative for edge density. Thus, area PM is positively modulated by contrast, but negatively modulated by edge density. (*p*-values are from one-way ANOVAs with the Tukey-Kramer correction for multiple comparison).

Neurons in the four imaged visual areas (V1, AL, LM, and PM) exhibited highly selective responses to the naturalistic video. Individual neurons responded to ∼ 3% of stimulus video frames, corresponding to a high lifetime sparseness (0.83 ± 0.09 (mean ± SD); **Fig. 2b**). Unbiased clustering (Gaussian mixture model, **GMM**) partitioned neurons into 25 tuning classes to the naturalistic video, and 20 of these classes exhibited unique sparse response patterns, responding at specific time points of the naturalistic video (**Supplementary Fig. 6d**). All naturalistic-video-tuning classes were observed in V1, and most were observed in HVAs (1-2 classes were missing in AL and PM). However, the relative abundance of tuning classes varied among V1 and HVAs (**Supplementary Fig. 6e, f**).

Next, we examined the collective effects of the biased distributions of tuning classes among HVAs. We reasoned that if a time-varying feature of the naturalistic video strongly modulates neuronal activity in an HVA (**Fig. 2c**), we should be able to detect that by regressing the visual feature dynamics (**Fig. 2d**) with the average neural activity in an HVA. The average response of a cortical area neuron population converged with several hundreds of neurons (about 500 from V1, about 200 from HVAs; **Supplementary Fig. 6c**). We examined a set of visual features that were previously implied to modulate visual system, including contrast^32,34^, luminance, edge density^26^, DOG entropy^33^, and OF speed and direction^23^ (**Supplementary Fig. 5**). The visual features were computed at multiple spatial scales, and qualitatively similar results were observed across a wide range of scales. Here we present representative results: edge density maps with a Gaussian kernel of 2.35° (full width at half maximum, **FWHM**), and DOG entropy maps with a Gaussian kernel of 11.75° (FWHM, inner kernel; the outer kernel is two-fold larger in FWHM) (**Fig. 2d**).

Regressing these visual features with the average neuronal responses per HVA suggested that V1, LM, AL and PM are distinctly modulated by naturalistic video features. We defined the modulation coefficients as the coefficients of the linear model, and modulation power of each feature as the variance of responses explained by the model (*i.e.,* the *r*^2^ of a linear fit to a particular visual feature and the neural activity; **Fig. 2e**). The feature modulation analysis suggested that the average responses of AL populations but not the other three areas was correlated with OF speed entropy (**Fig. 2f**). In PM, activity was correlated with contrast and edge density but not DOG entropy (**Fig. 2f**). In both V1 and LM, but neither AL nor PM, activity was correlated with both contrast and DOG entropy (**Fig. 2f**). These results suggested that AL activity represents motion components in the naturalistic video, while LM and PM activity represents spatial components in the same naturalistic video.

Next, we returned to the tuning class analysis (**Supplementary Fig. 6**), and determined whether the segregation of motion and spatial representations we observed was consistent with the biased distribution of tuning classes across HVAs. Tuning classes were indeed differentially modulated by contrast, DOG entropy, edge density and optical flow speed entropy of the naturalistic video (**Supplementary Fig. 7a**). As expected, we found that overrepresented tuning classes within an HVA could explain the superior representation of a feature. Similarly, the underrepresented tuning classes explained the inferior representation of a feature within an HVA (**Supplementary Fig. 7b**). Together, these results indicate that motion information and spatial information are differentially represented among HVAs due to the distribution of tuning classes between them. Neurons in AL provided superior representations of motion features in a naturalistic video, and neurons in LM and PM provided superior representations of spatial features in the same naturalistic video.

### DOG entropy can support texture family encoding

While V1 and LM also provided high fidelity representations of texture, PM did not (**Fig. 1d**). However, with the naturalistic video, all three areas were modulated by several spatial features. PM was distinct in that it was relatively well modulated by edge density and poorly modulated by DOG entropy, compared to V1 and LM. Thus, we hypothesized that DOG entropy could facilitate texture encoding. We generated DOG entropy and edge density feature maps for texture stimuli (**Fig. 1c**, **Supplementary Fig. 8a**). Then, we asked whether these feature maps were sufficient to discriminate texture images from different classes, while also being tolerant to differences among textures from the same class. We examined these questions by training a linear classifier to discriminate textures within and across texture classes using DOG entropy features or edge density features (*Methods*, **Supplementary Fig. 8b**). As expected, we found that DOG entropy indeed performed better for discriminating texture images by classes. The linear classifier using the DOG entropy feature successfully classified 83.3% of inter- and intra-class texture image pairs with 9 ± 4% miss-classification rate, while the classifier using the edge density feature classified 67% of these pairs with 12 ± 4% error rate. Thus, we concluded that the superior representation of texture by V1 and LM (compared to PM) could be due to their modulation by DOG entropy features.

### Gabor models exhibited biased feature representations

To this point, the evidence indicates a distributed representation of visual features among HVAs. Could these differences be due to subtle biases in preferred temporal or spatial frequencies? Or are they indicative of more fundamental differences in the underlying tuning of neurons in HVAs? To address these questions, we examined neuron models that would reproduce the diverse encoding functions we observed in mouse visual cortex. We simulated neurons using a base model of a linear-nonlinear (**LNL**) cascade with Gabor filter-based linear kernels (**Fig. 3a**; *Methods*). The LNL cascade with Gabor filter is a classic model for visual cortical neurons^35^. However, recent studies suggested that multiple Gabor kernels are required for predicting V1 neuron responses in mice^36^ and generating tolerances to rotation, translation, and scale^24,37^. Separately, dimensionality analysis suggested that normalization is critical for capturing the diverse response profiles of V1 neurons to naturalistic stimuli^38^. Inspired by these findings, we designed several variations of the base model for testing.

**Figure 3.**
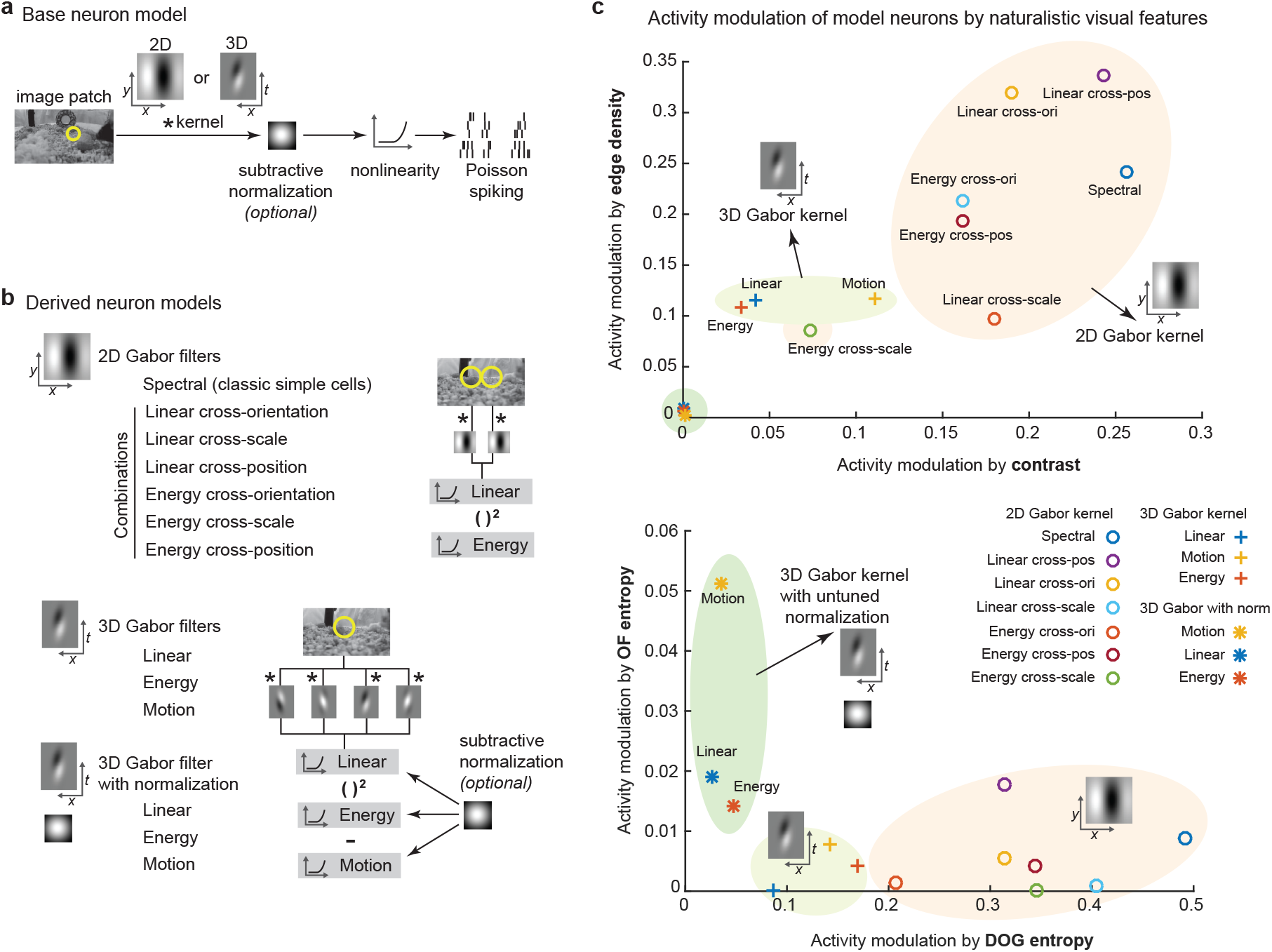
Spatial and motion feature encoding by variants of Gabor filter-based models. (**a**) The general architecture is a linear-nonlinear-Poisson (LNP) cascade neuron model. Neurons were simulated by various 2D and 3D Gabor-like linear kernels, with or without an untuned subtractive normalization. (**b**) From the base LNP model, variations were derived, organized into three classes: 2D Gabor-based, 3D Gabor-based without normalization, and 3D Gabor-based with normalization. Both linear and energy responses (akin to simple cells and complex cells) were computed from combinations of 2D Gabor filters. Linear, energy and motion responses (akin to simple cells, complex cells, and speed cells) were computed from 3D Gabor filters. (**c**) These three classes of models varied in how much their activity was modulated by global, form, and motion features in naturalistic videos. The neuron models are plotted by their modulation in feature spaces. The local of a neuron model was defined by the modulation power (same as Fig. 2e).

Models were grouped into three groups: 2D Gabor models, 3D Gabor models and 3D Gabor models with normalization. For 2D Gabor filter-based models, we examined both linear and energy models. These are similar to models of complex cells in which input from multiple simple cells with similar orientation preferences but varying phase are integrated^14^. Other combinations were used as well (cross-orientation, cross-scale, etc.; **Fig. 3b**). For 3D Gabor filter-based models, we also examined motion models (**Fig. 3b**). In addition, we also examined a version of the 3D Gabor model with subtractive normalization (**Fig. 3b**). All the simulations were carried out at multiple spatial and temporal (for 3D Gabor filter-based models) scales and sampled uniformly in space.

Using these three model classes (2D Gabor, 3D Gabor, and 3D Gabor with normalization), we simulated neuronal responses to the texture, RDK, and naturalistic video stimuli. We characterized the mutual information and feature selectivity of simulated responses to the texture and RDK stimuli (**Supplementary Fig. 9, 10**), and measured the feature encoding of simulated responses to the naturalistic video (**Supplementary Fig. 11**). Different neuron models varied in the encoding power of different types of stimuli or visual features. We noted that 2D Gabor models exhibited specific tuning to the texture family while remaining tolerant to motion directions, especially the cross-orientation and linear cross-position models (**Supplementary Fig. 9b**), which are the best models for texture family encoding. On the other hand, 2D Gabor models performed badly in representing the RDK stimuli (**Supplementary Fig. 10a**), while 3D Gabor models with normalization performed the best in encoding the RDK moving direction (**Supplementary Fig. 10a**). 3D Gabor models with untuned normalization captured both the information about the motion direction, but also exhibited tolerance to various coherence levels (**Supplementary Fig. 10b**). In representing the naturalistic videos, 2D Gabor models exhibited better sensitivity to the contrast, edge density, and the DOG entropy, while the 3D Gabor models with untuned normalization exhibited better modulation by the OF entropy (**Fig. 3c**). This represents an apparent trade-off in representation fidelity between 3D Gabor kernels with normalization and 2D Gabor kernels. In summary, the subtractive untuned normalization is important for the representation of motion, such as RDK and OF entropy, while Gabor kernels without the time domain provide better representations of spatial features.

### Gabor models reproduced specific feature representation of mouse visual cortex

With the model results in hand, we sought to determine how well they could account for our observations of neuronal activity *in vivo* (**Figs. 1,2**). We fit individual neuronal responses with the Gabor-based models (*Methods*). For each model class (2D Gabor, 3D Gabor, and 3D Gabor with normalization), one best linear model was fit by minimizing the cross-validation error of a linear regression between the simulated model response and neuron response (**Fig. 4a**).

**Figure 4.**
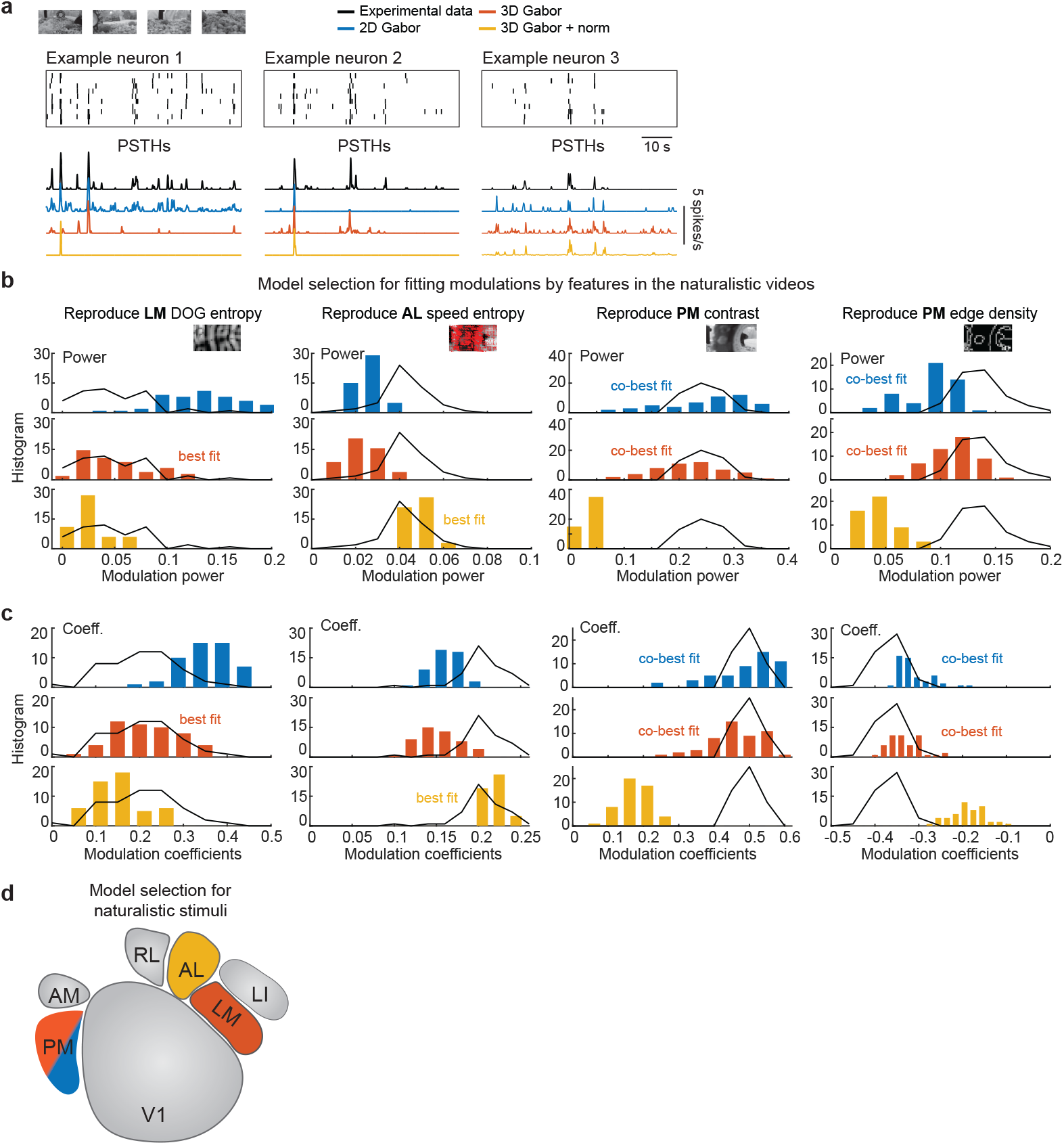
Segregated processing of spatial and motion features by visual neuron models. (**a**) Data from example neurons are shown in raster plots (top) and PSTHs (bottom), along with the best fits (as PSTHs) from each of the three model classes: 2D Gabor, 3D Gabor, and 3D Gabor with normalization. These three model fits, for each neuron, were used in the next analysis. (**b, c**) The three model neuron classes were characterized in terms of their modulation to global, form, and motion components of the naturalistic video. The distributions of (**b**) modulation power and (**c**) modulation coefficients for the three model classes were compared to those of the actual data, for neurons in specific HVAs and features those HVAs were well modulated by (see also, **Supplementary Fig. 7, 14**). The pool of model neurons for each pair of graphs (power and coefficient) for a cortical area were drawn from model fits to neurons in that same cortical area. (**d**) The diagram summarizes the model classes that best reproduce the modulation in three HVAs, LM, AL, and PM to global, form, and motion components in the naturalistic video.

Next, we took these pools of fits (three fits per neuron, one fit for each model class) and characterized how they represented features in the naturalistic videos. Consistent with the prior findings in this study (**Fig. 3**), we found that 2D Gabor models reproduced the neuronal information encoding about texture stimuli the best (**Supplementary Fig. 12**), while the 3D Gabor model with normalization reproduced the information encoding about the RDK stimuli the best (**Supplementary Fig. 13**).

The three classes of models were differentially involved in encoding of spatial and temporal features of naturalistic videos (**Supplementary Fig. 14**). We examined how well the three model classes could account for the characteristic neuronal activity modulations to visual features we observed in each HVA *in vivo*. We took a subset of the model responses, those that were fit to neurons that belonged to over-represented tuning classes within an HVA (**Supplementary Fig. 6d, f**), as these neurons accounted for unique spatiotemporal feature representations of HVAs (**Supplementary Fig. 7**). For the DOG entropy modulation in area LM, we found that the 3D Gabor model class best fit the modulation we observed *in vivo* (**Fig. 4b, c**). For the optic flow speed entropy modulation in area AL, model fits to AL neurons that were in the 3D Gabor with normalization class were the best fit to the *in vivo* data (**Fig. 4b, c**). For the contrast and edge density modulations observed in area PM, both 2D and 3D Gabor model classes provided good fits. However, the fits for the contrast modulation were a better match to the *in vivo* data than the edge density modulation (**Fig. 4b, c**). Overall, this analysis reveals that unique model classes are required to reproduce the visual feature modulation observed in HVAs: 3D Gabor filter-based models for area LM neurons, 3D Gabor filters with normalization for area AL neurons, and both 2D and 3D Gabor filter-based models for area PM neurons (**Fig. 4d**).

## Discussion

In the current study, we have revealed unique encoding properties of V1 and multiple HVAs in representing textures, RDK, and naturalistic videos. From our results, it appears as though V1 establishes a representation of various visual features, LM and LI are specialized for encoding of spatial features, and AL is specialized for the encoding of motion features. The encoding function of area PM was less obvious, as it seems that activity in that HVA was driven mostly by the density of visual edges, which are a spatial feature, but since PM is so poorly modulated by the DOG entropy feature, it is difficult to group it with LM and LI. Finally, we determined that unique model classes are required to reproduce the modulations we observed in these. Parameter variations within a model class were not sufficient. Instead, different model classes were required for reproducing the *in vivo* results in separate HVAs. These findings provide new insights into the neural circuitry that can generate distributed representations of visual stimuli in HVAs.

In our analysis, we found discrete neuron classes that had unique response profiles to a naturalistic video stimulus. These classes formed a non-uniform distribution among V1 and HVAs, and appropriately, were found to contribute to the biases in feature encoding among HVAs. It is unclear whether neurons with different tuning profiles play similar computational roles. Overall, these results determined that mouse visual cortical neurons can represent complementary features of visual scenes, and each HVA can exhibit unique biases towards specific visual features that are consistent across stimulus types, including naturalistic videos. Coupled with their downstream connectivity, these distinguishing biases among HVAs can provide insights into their involvement in visual processing and behavior.

The rodent visual system evolved in response to the ecological niche mice found themselves in. We do not expect such a process to result in neural circuitry that performs neat, absolute segregations of information about visual scenes. Instead, we expect neural circuitry that efficiently supports adaptive behavior for the mouse’s ecological niche. The principles of that efficient circuitry are likely quite different from those of any systematic, mathematically compact approach for parsing a visual scene in terms of known receptive field properties of visual cortical neurons. Thus, here we used a data-driven approach to gain a conservative foothold into complex visual scene processing in mice. We explored how segregated representations might emerge using a modeling approach based on known receptive field properties of visual cortical neurons, or at least popular models thereof. This analysis showed that 2D and 3D Gabor models provided accurate accounts for distinguishing texture and form features. By contrast, 3D Gabor models with subtractive normalization were key for distinguishing motion stimuli.

The enrichment of specific representations of motion or texture in areas AL and LM respectively, could arise from specific connectivity from other brain regions (e.g., V1) that preserves selectivity^10^, or from converging inputs that result in enhanced selectivity (or more invariant selectivity) for a visual feature^14,36,39^. We generally cannot distinguish those two possibilities with this data set. However, in area LI, neurons exhibited selectivity that surpassed that of neurons in V1, so it appears as though preserved selectivity from V1 projecting to LI would be insufficient to produce such selectivity. However, we cannot rule out thresholding effects which could play a role in increasing apparent selectivity.

Altogether, this study reveals new segregations of visual encoding or representations among HVAs in mice, many of which are reminiscent of the primate visual system. Studies have suggested macaque V2 exhibited selectivity to texture families and tolerance to local feature differences between images from the same texture class^24,26^. In macaque V4, neurons are highly selective to texture patterns, which are well predicted using combination of 2D Gabor models^37^. Famously, macaque dorsal visual areas such as MT exhibit selectivity to RDK motion direction^40^. The functional similarities between mouse LM and LI and macaque V2 and V4, and between mouse AL and macaque MT are perhaps superficial, but could also indicate that the dual stream framework for visual pathways in primates could have an analog in mice^5,41^. Earlier anatomical and receptive field mapping studies suggest that mouse LM and AL likely serve as the ventral and dorsal gateways in the mouse visual hierarchy^6,8,31^. Anatomical evidence including connectivity with downstream brain regions support functional distinctions between putative ventral and dorsal areas of mouse visual cortical areas, *e.g.* ventral areas were strongly connected to temporal and parahippocampal cortices, while putative dorsal areas were preferentially connected to parietal, motor and limbic areas^5^. Recent large scale multi-region electrode recordings from mouse visual cortex revealed an inter-area functional connectivity hierarchy, but did not group mouse HVAs into separate streams or subnetworks^9^. The study further showed that both LM and AL were similarly recruited by a visual recognition task, in which AM and PM were strongly involved^9^. Together, we conclude that both anatomical and functional studies suggest that mouse HVAs likely play distinct roles in visual behaviors, and may comprise dual processing streams analogous to primates. However, well designed behavioral tasks are required to further reveal the circuits and mechanisms.

## Methods

### Animal and surgery

All animal procedures and experiments were approved by the Institutional Animal Care and Use Committee of the University of North Carolina at Chapel Hill or the University of California Santa Barbara and performed in accordance with the regulation of the US Department of Health and Human Services. GCaMP6s-expressing transgenic adult mice of both sexes were used in this study. Mice were 110 – 300 days old for data collection. GCaMP6s-expressing mice were induced by triple crossing of the following mouse lines: TITL-GCaMP6s (Allen Institute Ai94), Emx1-Cre (Jackson Labs #005628), and ROSA:LNL:tTA (Jackson Labs #011008)^29^. Mice were housed under a 12 h light / 12 h dark cycle, and experiments were performed during the dark cycle of mice. For cranial window implantation, mice were anesthetized with isoflurane (1.5 – 1.8% in oxygen) and acepromazine (1.5 – 1.8 mg/kg body weight). Carprofen (5 mg/kg body weight) was administered prior to surgery. Body temperature was maintained using physically activated heat packs or homeothermic heat pads during surgery. Eyes were kept moist with ophthalmic ointment during surgery. The scalp overlaying the right visual cortex was removed, and a custom steel headplate with a 5 mm diameter opening was mounted to the skull with cyanoacrylate glue (Oasis Medical) and dental acrylic (Lang Dental). A 4 mm diameter craniotomy was performed over visual cortex and covered with a #1 thickness coverslip, which was secured with cyanoacrylate glue.

### Locating visual areas with intrinsic signal optical imaging (ISOI)

Prior to two-photon imaging, the locations of primary and higher visual area were mapped using ISOI, as previously reported^28,31,42^. Pial vasculature images and intrinsic signal images were collected using a CCD camera (Teledyne DALSA 1M30) and a tandem lens macroscope. A 4.7 × 4.7 mm^2^ cortical area was imaged at 9.2 μm/pixel spatial resolution and at 30 Hz frame rate. The pial vasculature was illuminated and captured through green filters (550 ± 50 nm and 560 ± 5 nm, Edmund Optics). The ISO images were collected after focusing 600 μm down into the brain from the pial surface. The intrinsic signals were illuminated and captured through red filters (700 ± 38 nm, Chroma and 700 ± 5 nm, Edmund Optics). Custom ISOI instrumentation were adapted from Kalatsky and Stryker^12^. Custom acquisition software for ISOI imaging collection was adapted from David Ferster^28^. During ISOI, mice were 20 cm from a flat monitor (60 × 34 cm^2^), which covered the visual field (110° x 75°) of the left eye. Mice were lightly anesthetized with isoflurane (0.5%) and acepromazine (1.5 – 3 mg/kg). The body temperature was maintained at 37 °C using a custom electric heat pad^28^. Intrinsic signal responses to vertical and horizontal drifting bars were used to generate retinotopic maps for azimuth and elevation. The retinotopic maps were then used to locate V1 and HVAs (**Supplementary Fig. 1a**). Borders between these areas were drawn using features of the elevation and azimuth retinotopic maps, such as reversals, manually^18,31^. The vasculature map provided landmarks to identify visual areas in two-photon imaging.

### *In vivo* two-photon imaging

Two-photon imaging was performed using a custom Trepan2p microscope controlled by custom LabView software^28^. Two regions were imaged simultaneously using temporal multiplexing^28^. Two-photon excitation light from an ultrafast Ti:Sapph laser tuned to 910 nm (MaiTai DeepSee; Newport Spectra-Physics) laser was split into two beams through polarization optics, and one path was delayed 6.25 ns relative to the other. The two beams were steered independently from each other using custom voice coil steering mirrors and tunable lenses. This way, the X, Y, Z plane of the two paths can be independently positioned anywhere in the full field (4.4 mm diameter). The two beams were raster scanned synchronously about their independently positioned centers by a 4 kHz resonant scanner and a linear scanner (Cambridge Technologies). Photons were detected (H7422P-40, Hamamatsu) and demultiplexed using fast electronics. For four-region scanning, the steering of the two beams was alternated every other frame.

In the current study, two-photon imaging of 500 x 500 μm^2^ was collected at 13.3 Hz for two-region imaging, or 6.67 Hz for quad-region imaging. We typically imaged neurons in V1 and one or more HVAs simultaneously. Up to 500 neurons (V1: 129 ± 92; HVAs: 94 ± 72; mean ± SD) were recorded per imaging region (500 x 500 μm^2^). Imaging was performed with typically <80 mW of 910 nm excitation light out of the front of the objective (0.45 NA), including both multiplexed beams together. Mice were head-fixed about 11 cm from a flat monitor, with their left eye facing the monitor, during imaging. The stimulus display monitor covered 70° x 45° the left visual field. Two-photon images were recorded from awake mice. During two-photon imaging, we monitored the pupil position and diameter using a custom-controlled CMOS camera (GigE, Omron) at 20 – 25 fps. No additional illumination was used for pupil imaging.

### Calcium imaging and imaging processing

Calcium imaging processing was carried out using custom MATLAB codes. Two-photon calcium imaging was motion corrected using Suite2p subpixel registration module^43^. Neuron ROIs and cellular calcium traces were extracted from imaging stacks using custom code adapted from Suit2p modules^43^. Neuropil contamination was corrected by subtracting the common time series (1^st^ principal component) of a spherical surrounding mask of each neuron from the cellular calcium traces^17,44^. Neuropil contamination corrected calcium traces were then deconvolved using a Markov chain Monte Carlo (**MCMC**) methods^44,45^. For each calcium trace, we repeated the MCMC simulation for 400 times, and measured the signal-to-noise of MCMC spike train inference for each cell (**Supplementary Fig. 1b**). Neurons in V1 and HVAs exhibited similar instantaneous firing rates (**Fig. 1c**). For all subsequent analysis, only cells that reliable spike train inference results were included (correlations between MCMC simulations is greater than 0.2).

### Visual stimuli

Visual stimuli were displayed on a 60 Hz LCD monitor (9.2 x 15 cm^2^). All stimuli were displayed in full contrast.

The texture stimuli (**Supplementary Fig. 2a**) were generated by panning a window over a large synthesized naturalistic texture image at one of the cardinal directions at the speed of 32 °/s. We generated the large texture image by matching the statistics of naturally occurring texture patterns^46^. The texture pattern families were: animal fur, mouse chow, rocks, and tree trunk. Each texture stimulus ran for 4 s and were interleaved by a 4 s gray screen.

The random dot kinematogram (**RDK**) stimuli contained a percentage (i.e., coherence) of white dots that move in the same direction (i.e., global motion direction) on a black background (**Supplementary Fig. 4a**). We presented the animal with RDK at three coherence levels (40%, 70%, and 90%) and four cardinal directions. The dot diameter was 3.8° and the dot speed was 48 °/s. White dots covered about 12.5% of the screen. The lifetime of individual dots were about 10 frames (1/6 s). These parameters were selected based on mouse behavior in a psychometric RDK task^47^. Each RDK stimulus ran for 3 – 7 s (responses in the first 3 s were used for analysis) and interleaved with 3 s gray screen. The same RDK pattern was looped over trials.

Two naturalistic videos (**Fig. 2a**) were taken by navigating a mouse home cage, with or without a mouse in the cage. Each video had a duration of 32 s and were presented with interleaved 8 s long periods with a gray screen. For the convenience of analysis, we concatenated the responses to the two videos (total 64 s).

### Visual features of the naturalistic video

We characterized various visual features of the naturalistic video (**Supplementary Fig. 5**).

*Average luminance*: The average pixel value of each frame.

*Global contrast*: The ratio between the standard deviation of pixel values in a frame, and the average luminance of that same frame.

*Edge density*: The local edges were detected by a Canny edge detector^48^. The algorithm finds edges by the local intensity gradient and guarantees to keep the maximum edge in a neighborhood while suppressing non-maximum edges. We applied the Canny edge detector after Gaussian blurring of the original image at multiple scales (1°-10°). A binary edge map was generated as the result of edge detection (**Supplementary Fig. 5a**). The edge density was computed as the sum of positive pixels in the binary edge map of each frame.

*Difference of Gaussian (**DOG**) entropy*: We characterized local luminance features following difference of Gaussian filtering at multiple scales, and then computed the entropy of these features within a local neighborhood (**Supplementary Fig. 5b**).

*Optical flow entropy*: We estimated the direction and speed of salient features (e.g., moving objects) using the Horn-Schunck method at multiple spatial scales. Then we computed the entropy of the OF direction and speed at each frame. Since the OF estimation relies on the saliency of visual features, the moving texture and RDK stimuli resulted in distinct OF entropies, with the latter being larger (**Supplementary Fig. 5c**).

Visual features were computed either by average over space or by computing a spatial variance value (i.e. entropy). These measurements were inspired by the efficient coding theory^49^, which suggested that the neuron population coding is related to the abundance or the variance of visual features in the natural environment.

### Reliability and sparseness

The reliability of responses to naturalistic videos was defined as the trial-to-trial Pearson correlation between inferred spike trains of each neuron binned in 500 ms bins. The reliability of responses to texture stimuli and RDK were computed as the fraction of trials that a neuron fired to its preferred stimulus within a time window (2 s for texture stimuli and 3 s for RDK). These definitions were commonly used in previous studies^39,50^. Only reliably responsive neurons were included in the latter analysis (Pearson correlation > 0.08 to naturalistic video; fired on > 60% trials to the texture and RDK stimuli). The qualitative results were not acutely sensitive to the selection criteria.

The sparseness was computed as (eq. 1)^51^:

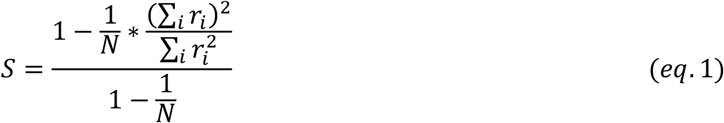

For lifetime sparseness, *r_i_* is trial-averaged response to i^th^ stimulus and N is the length of the stimuli. The sparseness to naturalistic videos was computed using 500 ms bins. The qualitative results of reliability and sparseness were not acutely sensitive to the bin size.

### Gaussian mixture model

To characterize the tuning properties in an unbiased manner, neurons were clustered using a Gaussian mixture model^52^ (**GMM**) based on the trial-averaged responses to the naturalistic video. Only reliably responsive neurons were included for GMM analysis (trial-to-trial Pearson correlation of the inferred spike trains > 0.08, after spike trains were binned in 500 ms bins). Neuronal responses of the whole population, pooled over all cortical areas imaged, were first denoised and reduced in dimension by minimizing the prediction error of the trial-averaged response using principle component (PC) analysis. 55 PCs were kept for population responses to the naturalistic videos. We also tested a wide range of PCs (20 – 70) to see how this parameter affected clustering, and we found that the tuning group clustering was not acutely affected by the number of PCs used. Neurons collected from different visual areas and different animals were pooled together in training the GMM (3527 neurons). GMMs were trained using the MATLAB build function *fitgmdist* with a range of numbers of clusters. A model of 25 classes was selected based on the Bayesian information criterion (BIC). We also examined models with different numbers of classes (20, 30, 45, or 75), and found that the main results held regardless of the number of GMM classes. Neurons with similar response patterns were clustered into the same class. **Supplementary Fig. 6** shows the response pattern of GMM classes to the naturalistic video. The size of the naturalistic video classes are shown in **Supplementary Fig. 6d**. To examine the reproducibility of the GMM classification, we performed GMM clustering on 10 random subsets of neurons (90% of all neurons). We found the center of the Gaussian profile of each class was consistent (Pearson correlation of class centers, 0.74 +/- 0.12). About 65% of all neurons were correctly (based on the full data set) classified, while 72% of neurons in classes that are over-represented in HVAs were correctly classified. Among misclassifications, about 78% were due to confusion between the three untuned classes with tuned classes. Thus, most of the classes to come out of the GMM analysis appear to be reproducible, and are not sensitive to specific subsets of the data.

### Information analysis

Mutual information (**MI**) evaluates the information the neuronal response (*r*) has about certain aspects of the stimulus, and it is computed in units of bits. It was computed using the following equation.

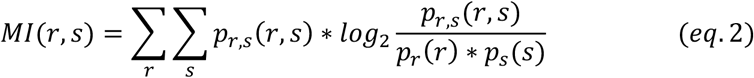

We computed the MI between neuron responses and the visual stimulus (*s* has 16 categories for texture stimuli, *p_s_*(*s*) = 1/16; *s* has 12 categories for RDK, *p_s_*(*s*) = 1/12). We also computed the MI between neuron responses and the texture family (*s* has 4 categories for texture stimuli, *p_s_*(*s*) = 1/4), and the MI between neuron responses and the moving directions (*s* has 4 categories for both texture stimuli and RDK, *p_s_*(*s*) = 1/4). The probability of neuron responses were computed from spike count distributions within a stimulus window (2 s for texture stimulus and 3 s for RDK). Reliable RDK and texture responsive neurons (reliability > 0.6), which fired for more than 60% of the trials to the preferred stimulus, were included for the MI analysis.

### Regularized encoder model

To estimated the encoding pattern of texture responsive neurons and RDK responsive neurons, i.e. which texture pattern one neuron responded to, or how many texture patterns one neuron responded to, we decomposed the neuronal responses into motion direction components, and texture family or RDK coherence components using singular value decomposition (**SVD**). To be more robust, instead of using trial-averaged response, we first estimated the neuronal responses by linearly regressing with a unit encoding space (**Supplementary Fig. 2c-e, 4c-d**). Lasso regularization was applied to minimize overfitting. The regularization hyper-parameters were selected by minimizing the cross-validation error in predicting single trial neuronal responses. The linear regression model performance was measured by the Pearson correlation between the trial-averaged neuron response and the model. Only well-fit neurons were included for the following analysis (model performance > 0.6; about 70% of the whole population). The model selection criteria did not affect the qualitative results.

We then characterized the SVD components of well-fit neurons. Well-fit neurons exhibited either zero, one, or multiple significant SVD components (eigenvalue > 1). Neurons with zero significant SVD component were untuned neurons, while neurons with multiple significant SVD components suggested complicated tuning properties. We went on to characterize neurons which had single significant SVD components, as for which the neuronal responses were decomposed into a motion directions vectors, and a texture pattern vector or a motion coherence vector unambiguously (**Supplementary Fig. 2d, 4c**).

About 40-70% of well-fit texture neurons and about 50 – 60% of well-fit RDK neurons had only one significant SVD component. We define positive motion directions, or texture patterns for each neuron, when its corresponding vector value (singular vector of SVD) is greater than 0.2 (for texture responses) or 0.3 (for RDK responses) (the threshold value did not affect qualitative results; **Supplementary Fig. 2d, 4c**). In the results section, we report the distributions for neurons with different numbers of positive motion directions, texture patterns, and coherence levels for HVAs (**Supplementary Fig. 2f, 4e**).

### Modulation power of naturalistic visual features

For each cortical area, neuronal activity in response to the video was pooled and averaged, after binning into 500 ms bins. Then, separately for each cortical area, a linear regression model was fit to the average population response with individual features. These features are described above in the section (*Visual features of the naturalistic video*). We then evaluated a feature’s contribution in modulating the average population responses by the variance explained (r-squared) of each model (**Fig. 2d, f**). Features were computed over multiple spatial scales. The spatial scales that best modulated (highest r-squared) the neuronal response was used for this analysis.

To evaluate the significance of neuron classes, we repeated this process using different source data. Instead of using a pool of neurons from a cortical area, we used a pool of neurons from a specific class (200 neurons per pool with permutation). Again, we averaged activity over the pool, and then determined which features modulated activity of the class (**Supplementary Fig. 7a**). This process was repeated for classes that were either over-represented in an HVA or under-represented in an HVA (**Supplementary Fig. 7b**).

### SVM discrimination of texture images

We computed the pairwise distance between texture images (**Supplementary Fig. 8a**) within the same class or from different classes (**Supplementary Fig. 8b**). The Euclidean distance was computed using of DOG entropy (11.75° spatial filter size) or edge density (2.35° spatial filter size) feature maps. We then trained a support vector machine (**SVM**) classifier to discriminate texture images within and across classes, based on this pairwise distance (using the Matlab built-in function *classify*). We reported the cross-validation classification error rate (**Supplementary Fig. 8b**).

### Simulation of Gabor-based models

The neuron models used the structure of a linear-nonlinear (**LNL**) cascade. The spiking of model neurons was simulated following a nonhomogeneous Poisson process with a time varying Poisson rate. The rate was calculated by convolving visual stimuli with a linear kernel or a combination of linear kernels, followed by an exponential nonlinearity (**Fig. 3a**). Linear kernels were modeled by 2D (XY spatial) or 3D Gabor (XYT spatiotemporal) filters defined over a wide range of spatiotemporal frequencies and orientations. We simulated neurons with simple cells, complex cells and speed cells models^53^ (**Fig. 3b**). The three differed in the linear components of the LNL cascade: simple cells (called *linear model*, or *spectral model* for the 2D Gabor kernels) used the linear response of a Gabor filter; complex cells (called *energy model*) used the sum of the squared responses from a quadrature pair of Gabor filters (90° phase shifted Gabor filter pairs); speed cells (called *motion model*) used the arithmetic difference between the energy responses from an opponent pair of complex cells. We also modeled neurons based on the cross product of the linear or energy responses from two 2D Gabor filters (called *combination model*). In particular, we simulated the following three combination models: 1. 2D Gabor filters matched in spatial scale and location but tuned to different orientations (*cross-orientation model*); 2. 2D Gabor filters tuned to the same orientation and location with different spatial scales (*cross-scale model*); 3. 2D Gabor filters with matched tuning properties but offset in visual space (*cross-position model*) (**Fig. 3b**). In addition, we included a subtractive normalization before taking the nonlinearity in some models. A total of 13 neuron model types were used (**Fig. 3b**).

To examine feature encoding by these neuron model types, we performed 10 – 20 repeats of simulation for each neuron model to each stimulus. Either the simulated spike trains or peristimulus time histograms (**PSTH**) were used for characterizing the feature encoding. We analyzed the model responses in the same way as we had done for the mouse experimental data. We computed the mutual information between simulated neuron responses and texture stimuli or RDK stimuli, and characterized the selectivity of simulated neurons to texture families or RDK directions (**Supplementary Fig. 9, 10**). Next, we examined the modulation of simulated population responses by visual features of the naturalistic video. Neuron models were located in the feature space by how much of the population response variance was explained by individual features (**Supplementary Fig. 11**).

### Reproducing neuron responses to stimuli with Gabor-based models

To reproduce the feature representation of HVAs with neuron models, we fit individual neuronal responses with models following a linear regression equation (eq. 3). The linear coefficients were optimized by minimizing the cross-validation error. We also tested a sigmoidal nonlinear fitting (eq. 4). Sigmoidal parameters were optimized through gradient descent. As sigmoidal nonlinearity did not significantly improve the modeling performance, we reported the results from the linear fitting.

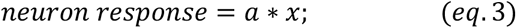

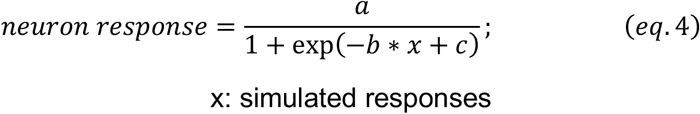

Neuron models were grouped into three categories: 2D Gabor models, 3D Gabor models, and 3D Gabor models with normalization. One model of each category, which minimize the cross-validation error, was selected for each neuron. The feature representation was then characterized on the model neuron responses.

### Data availability

All source data generating main figures will be available online upon publishing. All raw data are available upon request.

## Author contributions

All experiments and analysis were performed by YY. The imaging system was built by JNS. Animal handling was assisted by CRD. Study design and supervision by SLS. YY and SLS wrote the paper.

## Acknowledgements

Funding was provided by grants from the NIH (R01EY024294, R01NS091335), the NSF (1707287, 1450824), the Simons Foundation (SCGB325407), and the McKnight Foundation to SLS; a Helen Lyng White Fellowship to YY; a career award from Burroughs Welcome to JNS; and training grant support for CRD (T32NS007431).

## Supplementary Figures

**Supplementary Figure 1.**
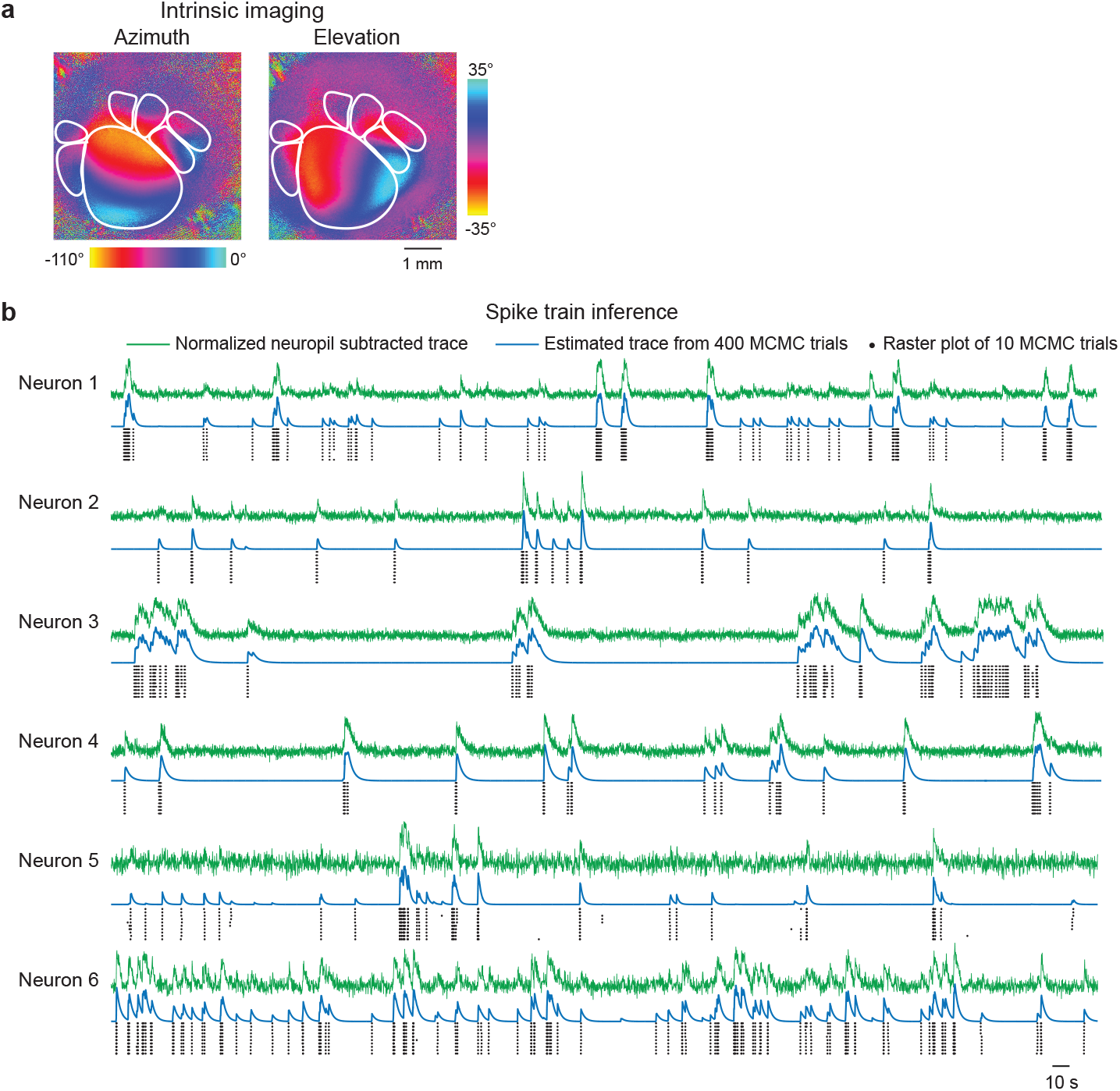
Multi-region two-photon calcium imaging processing. (**a**) Example intrinsic signal imaging of mouse visual areas. (**b**) Spike train inference of example neurons by Markov chain Monte Carlo (MCMC) methods.

**Supplementary Figure 2.**
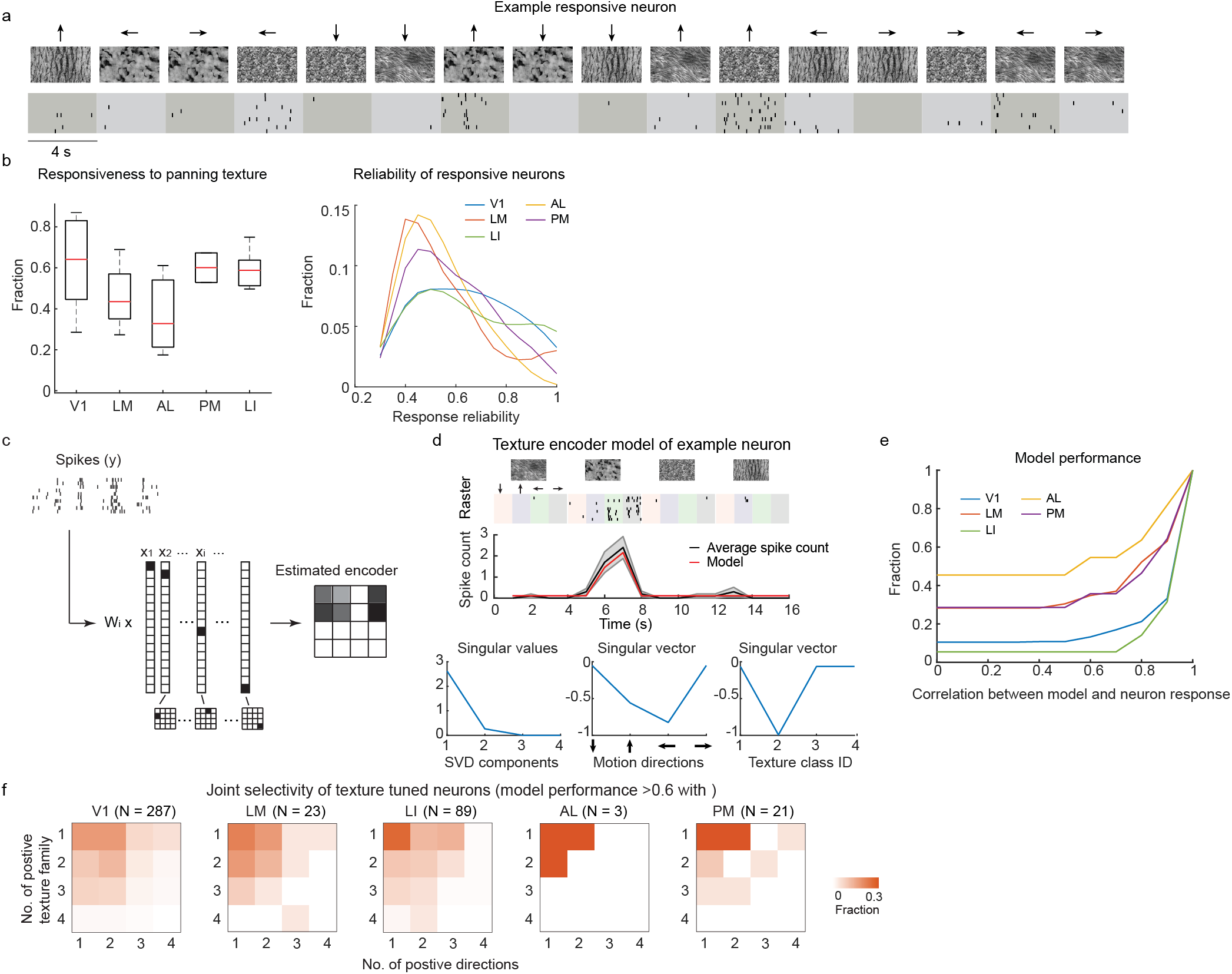
Defining tuning groups for neuronal responses to texture stimuli. (**a**) Example raster of a responsive neurons to moving texture stimuli with four texture families and four moving directions. (**b**) The responsiveness of V1 and HVAs to the texture stimuli (responsive neuron fires on more than 30% of the trials to the preferred stimulus). Left: the fraction of responsive neurons in HVAs are not significantly differed (one-way ANOVA, p = 0.2). Right: distribution of neuron firing reliability (firing probability over trials) to the preferred texture stimulus. Only responsive neuron was considered. V1 and LI were more reliable to the texture stimuli (one-way ANOVA with Bonferroni multiple comparison, p = 4 x 10^-10^). (**c**) Fit neuronal response (spike count) to an encoder model using least-square regression with lasso regularization. (**d**) Model performance of an example neurons. Top: raster plot and average spike count of the example neuron, overlaid with the estimated spike count from the model. The model spike count was highly correlated with the average spike count of the example neuron (Pearson correlation, r = 0.98). Bottom: SVD decomposition of the estimated encoder model. The left and right singular vectors corresponding to the motion direction and the texture family components, respectively. (**e**) Cumulative fraction of encoder model performance, which was defined as the Pearson correlation between model spike count and the trail-averaged spike count of neurons. (**f**) Joint distribution of the number of texture families and the number of directions that a texture neuron encoder was responsive to. Color hue indicate the fraction of neurons in each bin.

**Supplementary Figure 3.**
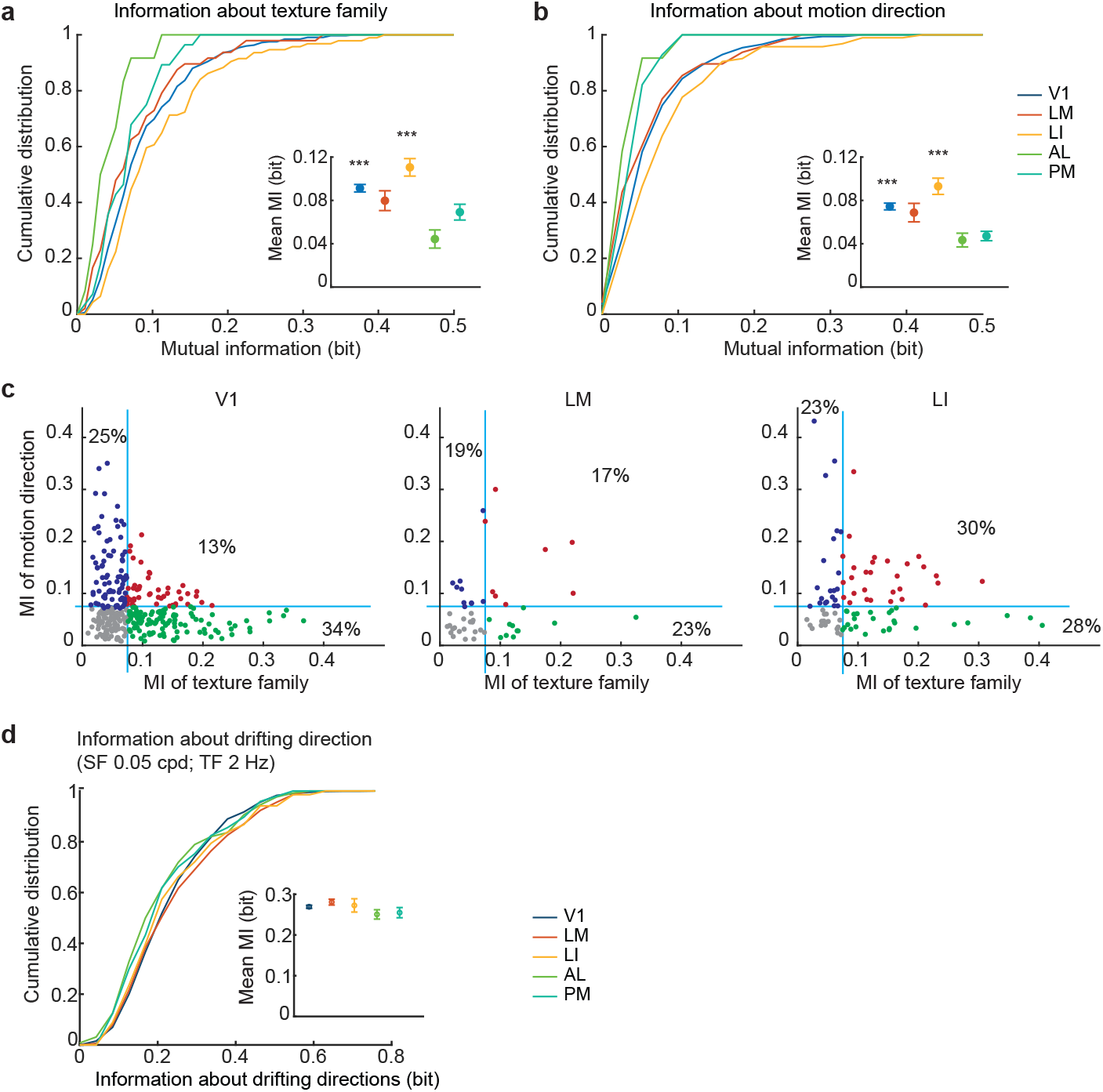
Information about texture stimuli is strongest in area LI. (**a-b**) Cumulative distribution of information each neuron has about the texture family (**a**) and the moving direction (**b**). Inserts are the mean information. Error bars indicate SE. V1 and LI carries significantly more information about the texture family (p = 0.0006) and the moving direction (p = 0.0006) (one-way ANOVA, Bonferroni multiple comparison). (**a-b**) Neurons reliably response (response to >60% of trials) to at least one texture stimuli were included for information analysis and encoder modeling (Number of neurons included (No. of experiments): V1, 325 (11); LM, 48 (5); LI, 96 (5); AL, 12 (3); PM, 28 (2)). (**c**) Relation between information about the moving direction and information about the texture family carried by individual neuron in V1, LM and LI. Each dot indicates one neuron. Blue line indicates the threshold of significant amount of information, which was defined by shuffled data (Mean + 3*SD). (**d**) Information about drifting grating directions were not striking differed among between HVAs (p = 0.12; one-way ANOVA).

**Supplementary Figure 4.**
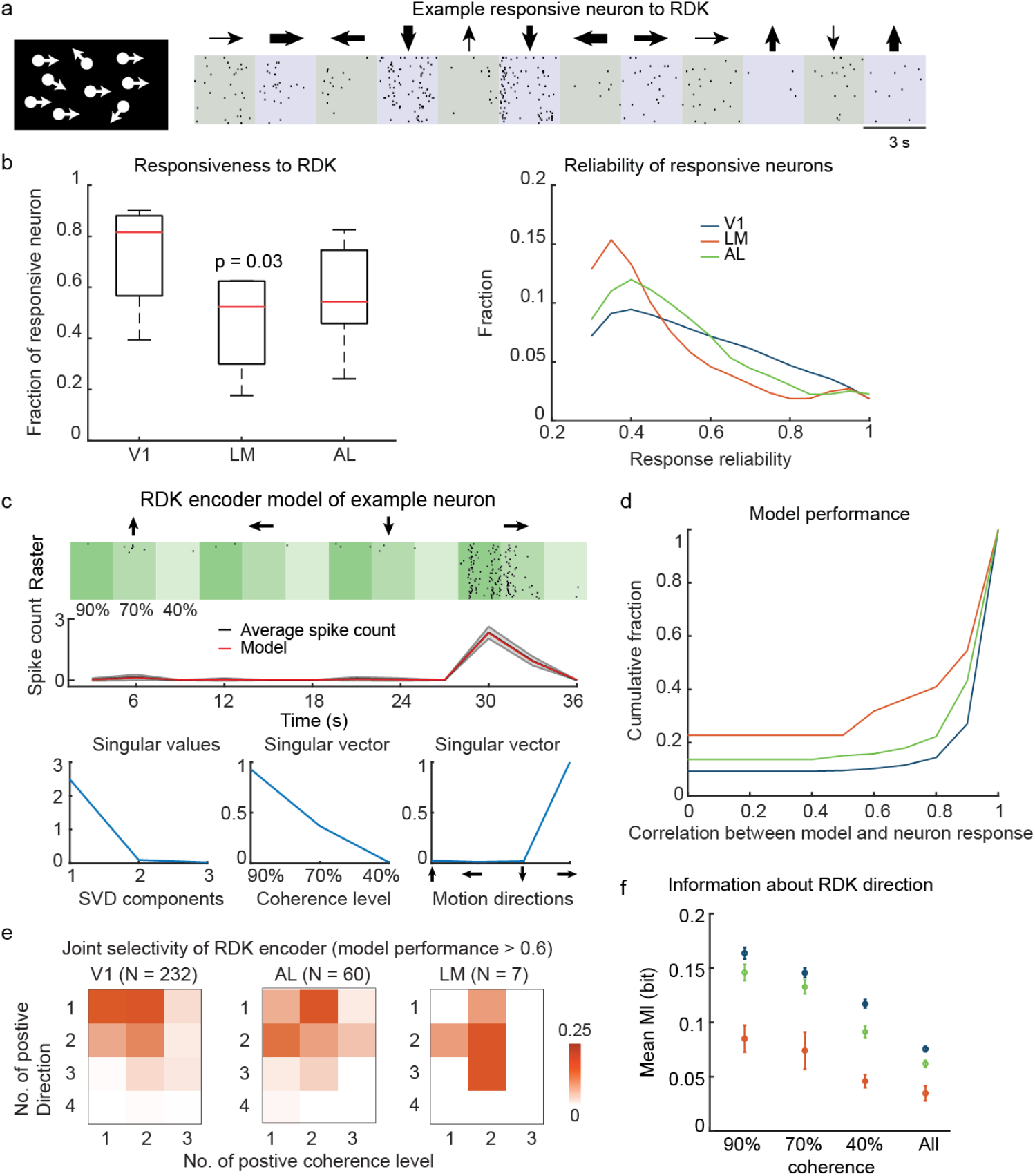
Defining tuning groups of neuronal responses to RDK stimuli. (**a**) Raster plots of an example neuron to RDK at four global motion directions and three coherence (indicated by the thickness of arrows). (**b**) The responsiveness of V1 and HVAs to the RDK stimuli (responsive neuron fires on more than 30% of the trials to the preferred stimulus). Left: the fraction of responsive neurons in LM was significantly smaller compare to V1 and AL (T-test, p = 0.03). Right: distribution of neuron firing reliability (firing probability on multiple trials) to the preferred RDK stimulus. Only responsive neuron was considered. V1 and AL were more reliable to the texture stimuli (one-way ANOVA with Bonferroni multiple comparison, p = 3 x 10^-5^). (**c**) RDK encoder model performance of an example neurons. Top: raster plot and average spike count of the example neuron, overlaid with the estimated spike count from the model (Pearson correlation, r = 0.99). Bottom: SVD decomposition of the estimated encoder model. The left and right singular vectors corresponding to the coherence level and the motion direction components, respectively. (**d**) Cumulative fraction of encoder model performance (Pearson correlation between model and the trail-averaged spike count of neurons). (**e**) Joint distribution of the number of directions and the number of coherence levels that an RDK neuron encoder was responsive to. V1 and AL has larger fraction of tuned neurons that were selectively response to one motion direction (V1, 52%; AL, 43% and LM, 14%). Color hue indicate the fraction of neurons in each bin. (**f**) The mean information about global moving direction at different coherence level.

**Supplementary Figure 5.**
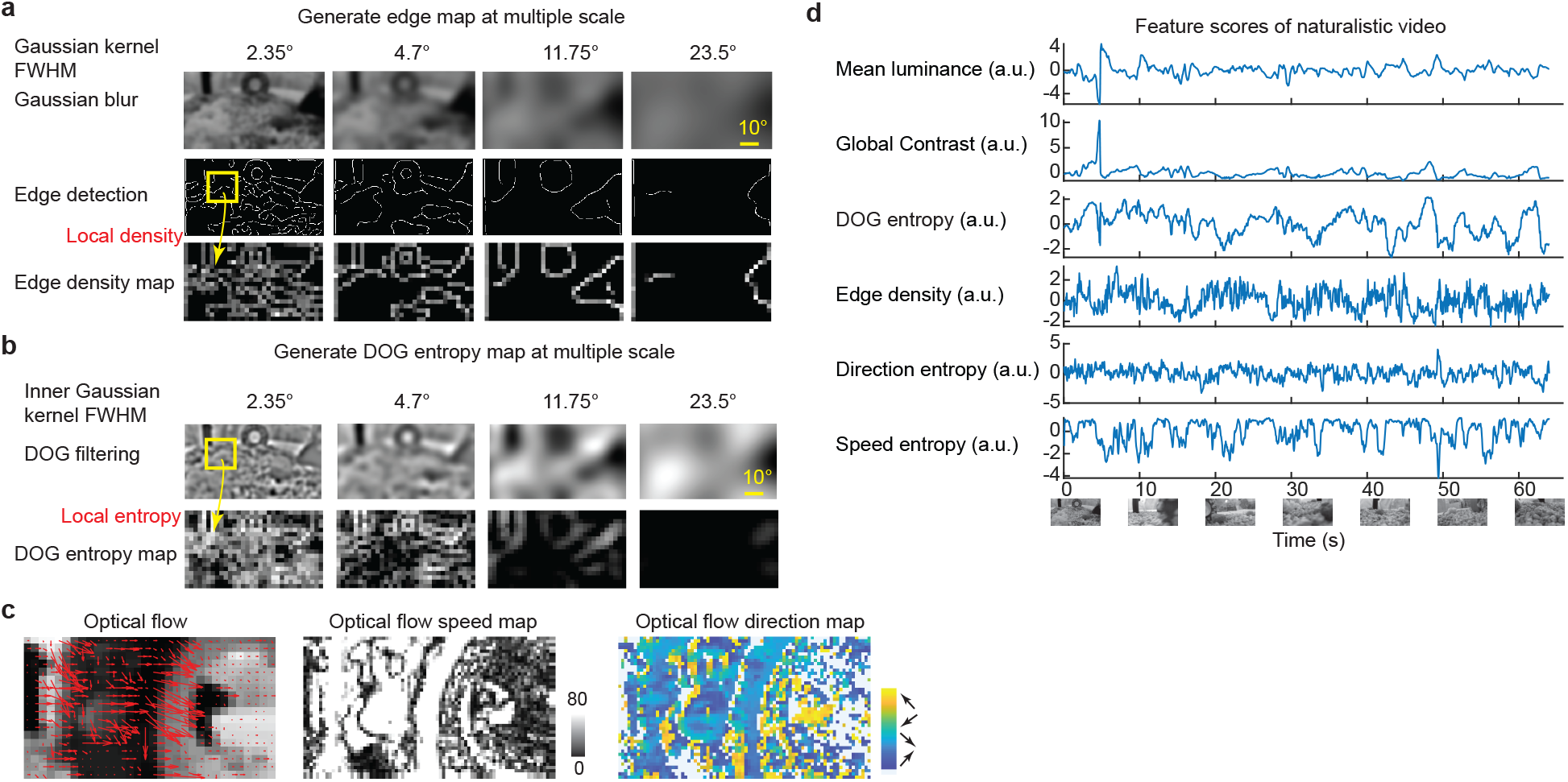
Feature space of naturalistic videos. (**a**) Example edge density maps at multiple spatial scales. Edge detection by Canny edge detector after Gaussian blur with defined kernel size (top). Edge density was computed by sum up the edge number within a local neighborhood (10 x 10 pixel, a wide range (10∼100 pixel^2^) of neighborhood size was tested). (**b**) Example Difference of Gaussian (**DOG**) entropy maps at multiple spatial scales. The inner Gaussian kernel size was shown (top), and outer Gaussian filter size was double the inner filter size. The entropy after DOG filtering was computed at a local neighborhood (10 x 10 pixel, a wide range (10∼100 pixel^2^) of neighborhood size was tested). (**c**) Example optical flow map for a naturalistic video frame. The OF direction and speed of each pixel was estimated using Horn-Schunck method. The OF feature entropy was computed at a local neighborhood. (**d**) The time-varying visual features of the naturalistic videos.

**Supplementary Figure 6.**
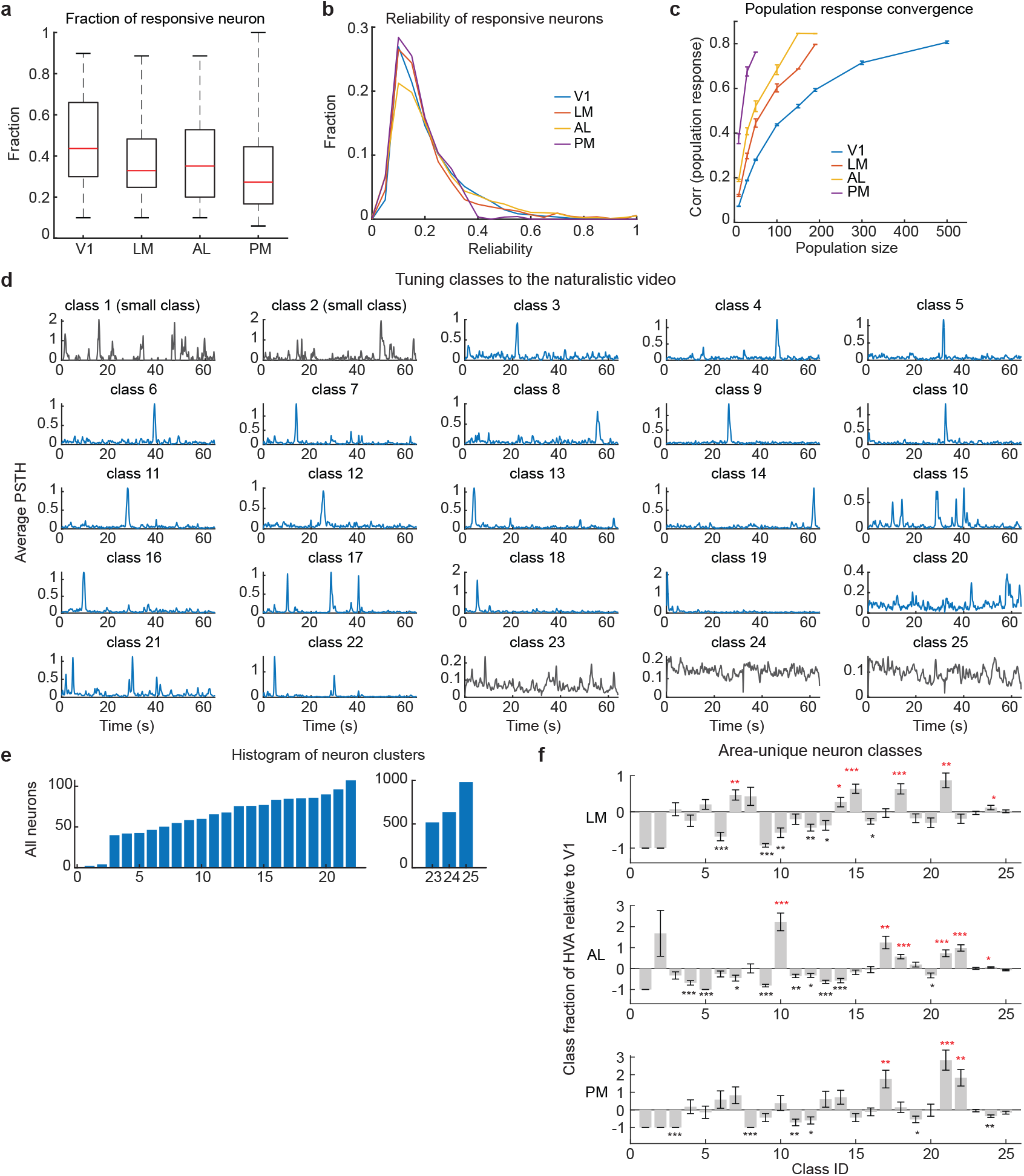
Selective response to a naturalistic video reveal classes that are enriched in specific HVAs. (**a**) The fraction of responsive neurons to the naturalistic video in V1, LM, AL and PM were similar (trial-to-trial Pearson correlation > 0.08; one-way ANOVA, p = 0.13). (**b**) The distribution of neuron firing reliability (trial-to-trial Pearson correlation) to the naturalistic video were not differed in V1, LM and AL, and slightly lower in PM (one-way ANOVA with Bonferroni multiple comparison, p = 0.0006). Only responsive neuron was considered. (**c**) Pearson correlation of the average population responses computed from non-overlapping subpopulations with certain number of neurons. (**d**) Average responses of 25 GMM classes to the naturalistic video. (**e**) Number of neurons in each GMM class to the naturalistic video. (**f**) Fraction difference of classes between HVA and V1 ( (HVA – V1) / V1). Zero means the HVA and V1 weighted the same on this class. Stars indicate significantly 10% more (red) or less (black) than V1 (T-test). The error bars indicate SE computed from permutation.

**Supplementary Figure 7.**
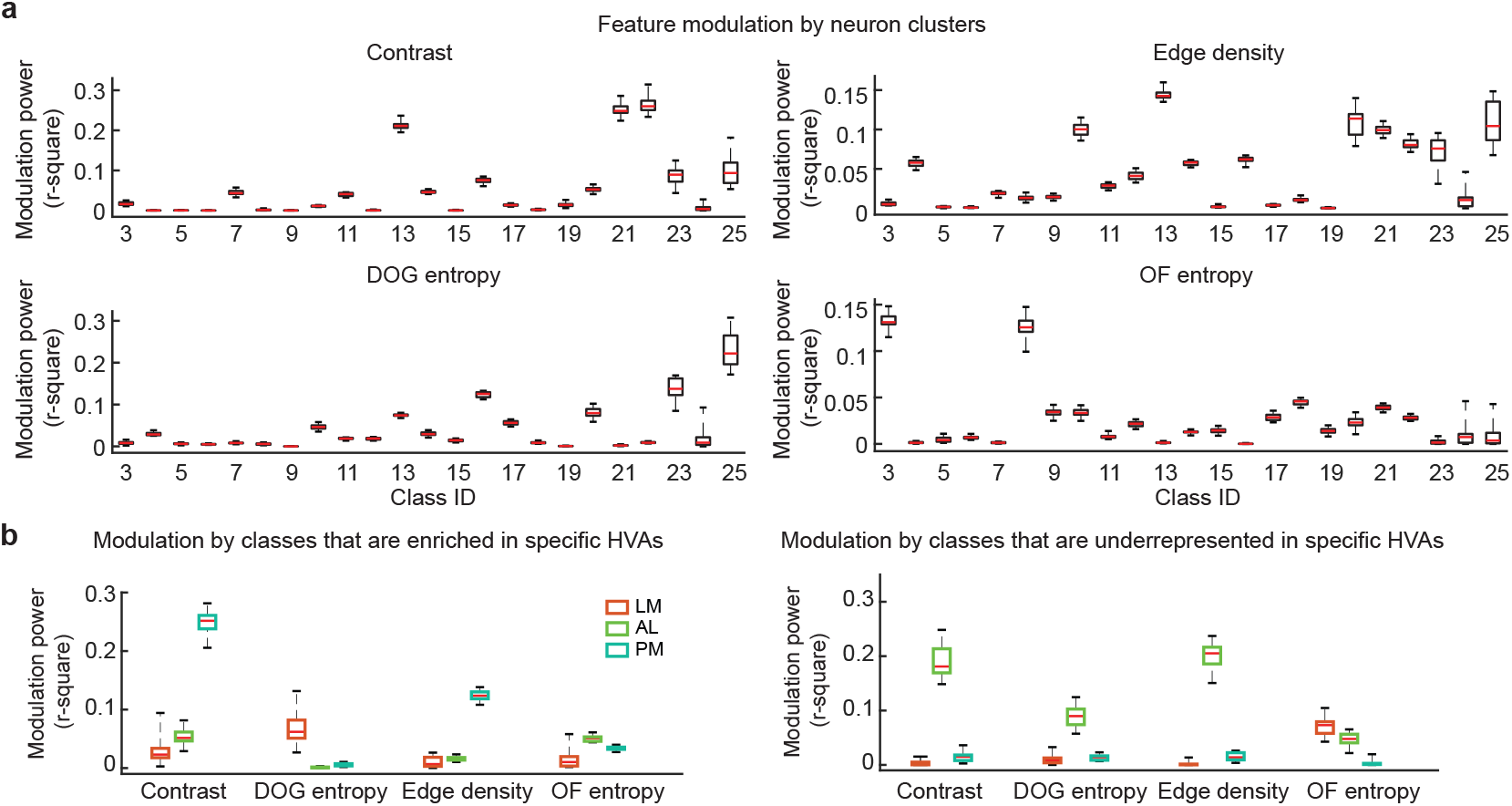
Parametric features of naturalistic video stimuli differentially modulate activity in the tuning groups. (**a**) The modulation power of the average responses of neurons in each class (N = 200 with permutation) by visual features of the naturalistic video. The modulation power is characterized by the r-squared values (variance explained) of the linear regression of the average population responses with individual features. (**b**) The modulation power of the average responses of a neuron population from selected classes (N = 200 with permutation) by visual features of the naturalistic video. Left, classes that are ENRICHED (more common than average of all HVAs) in specific HVAs (classes with red star in **Supplementary figure 6f**). Right, Classes that are UNDERREPRESENTED in specific HVAs (classes with black star in **Supplementary figure 6f**).

**Supplementary Figure 8.**
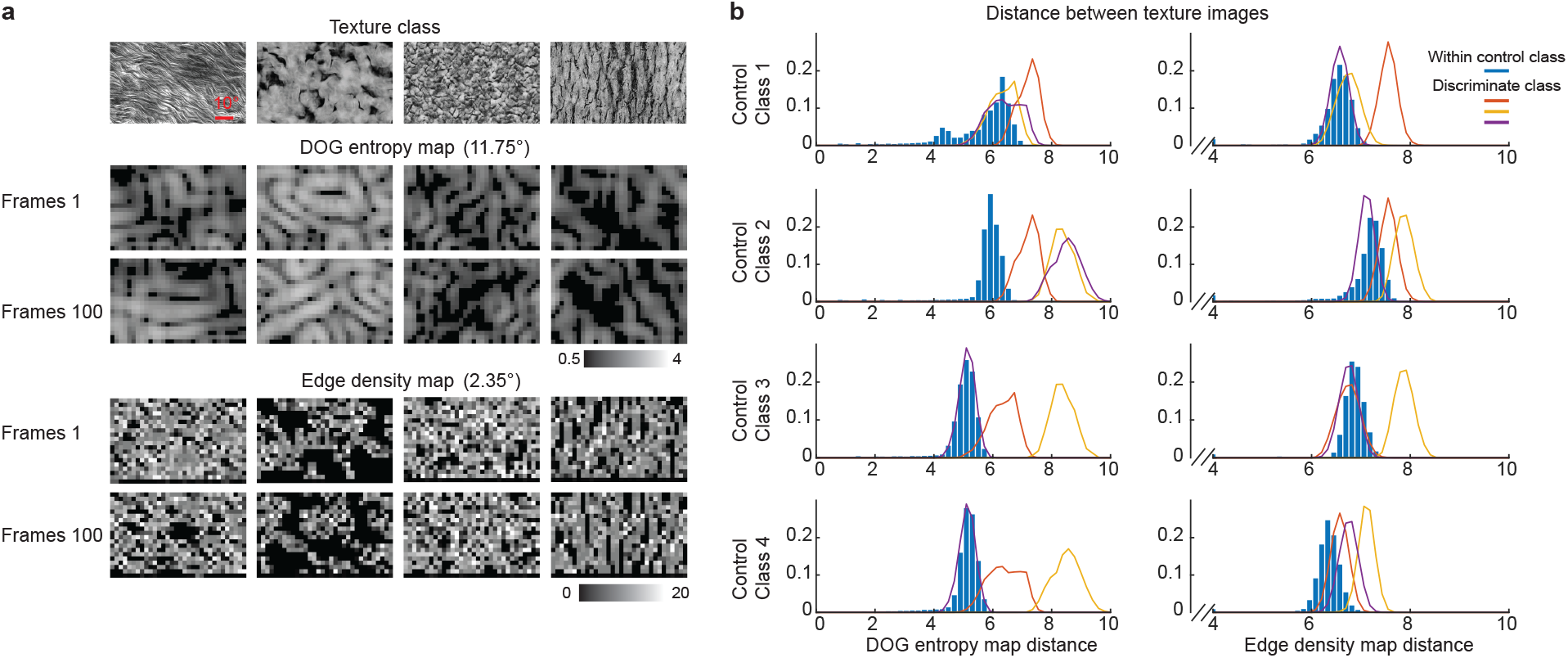
DOG entropy facilitated texture family encoding. (**a**) DOG entropy map and edge density map for example images from four texture classes. The Gaussian kernel size (standard deviation) is indicated in degrees. (**b**) Histogram of pairwise distances between texture images, from the same (blue bars) or different classes (colored curves). The distance was computed from the Euclidian distance between DOG entropy maps (left) or the edge density maps (right).

**Supplementary Figure 9.**
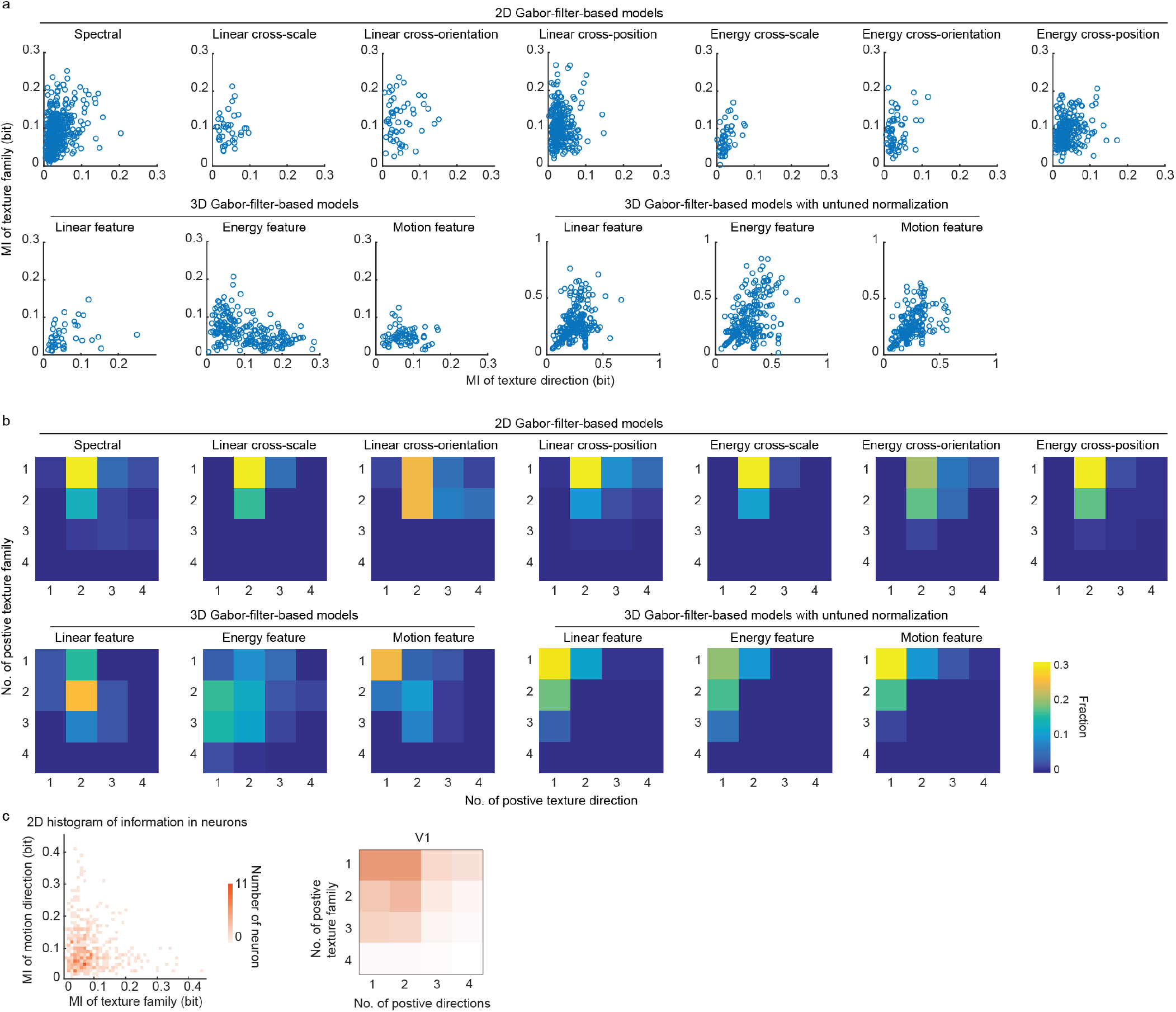
Encoding of texture stimuli by neuron models. (**a**) Plots of information about the texture family and information about the texture motion direction. Open circle indicates individual simulated neurons. 2D Gabor and 3D Gabor models without normalization exhibited unimodal encoding of the moving texture stimuli, that the former carried information about texture family and the latter carried information about texture motion direction. 3D Gabor models with untuned normalization generated joint encoding of both the texture family and the texture motion direction. (**b**) Joint distribution of the number of texture families and the number of texture moving directions that a simulated neuron was responsive to. (**c**) Left, the joint distribution of information about texture family and texture direction of mouse visual cortical neurons (reproduce **Supplementary Fig. 3c**, combined all regions). Right, Joint selectivity of texture families and texture moving directions mouse visual cortical responses (reproduce **Supplementary Fig. 2f**).

**Supplementary Figure 10.**
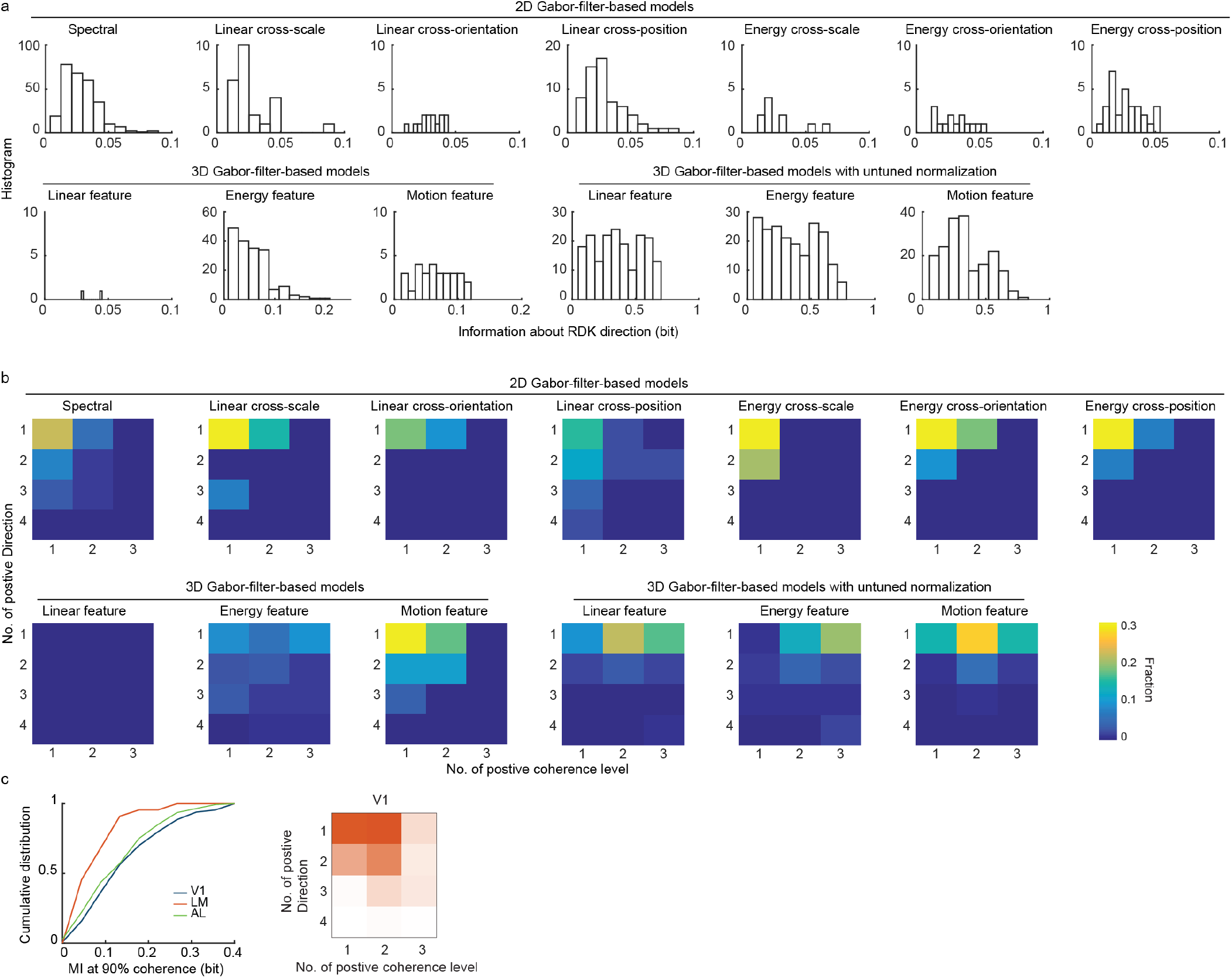
Encoding of RDK stimuli by neuron models. (**a**) Histogram of information about the RDK moving direction. (**b**) Joint distribution of the number of directions and the number of coherence levels that a simulated neuron was responsive to. (**c**) Left, the distribution of information about RDK motion direction of visual cortical neurons (reproduce Fig. 1f). Right, the selectivity of RDK directions of mouse visual cortical neurons (reproduce **Supplementary Fig. 4e**)

**Supplementary Figure 11.**
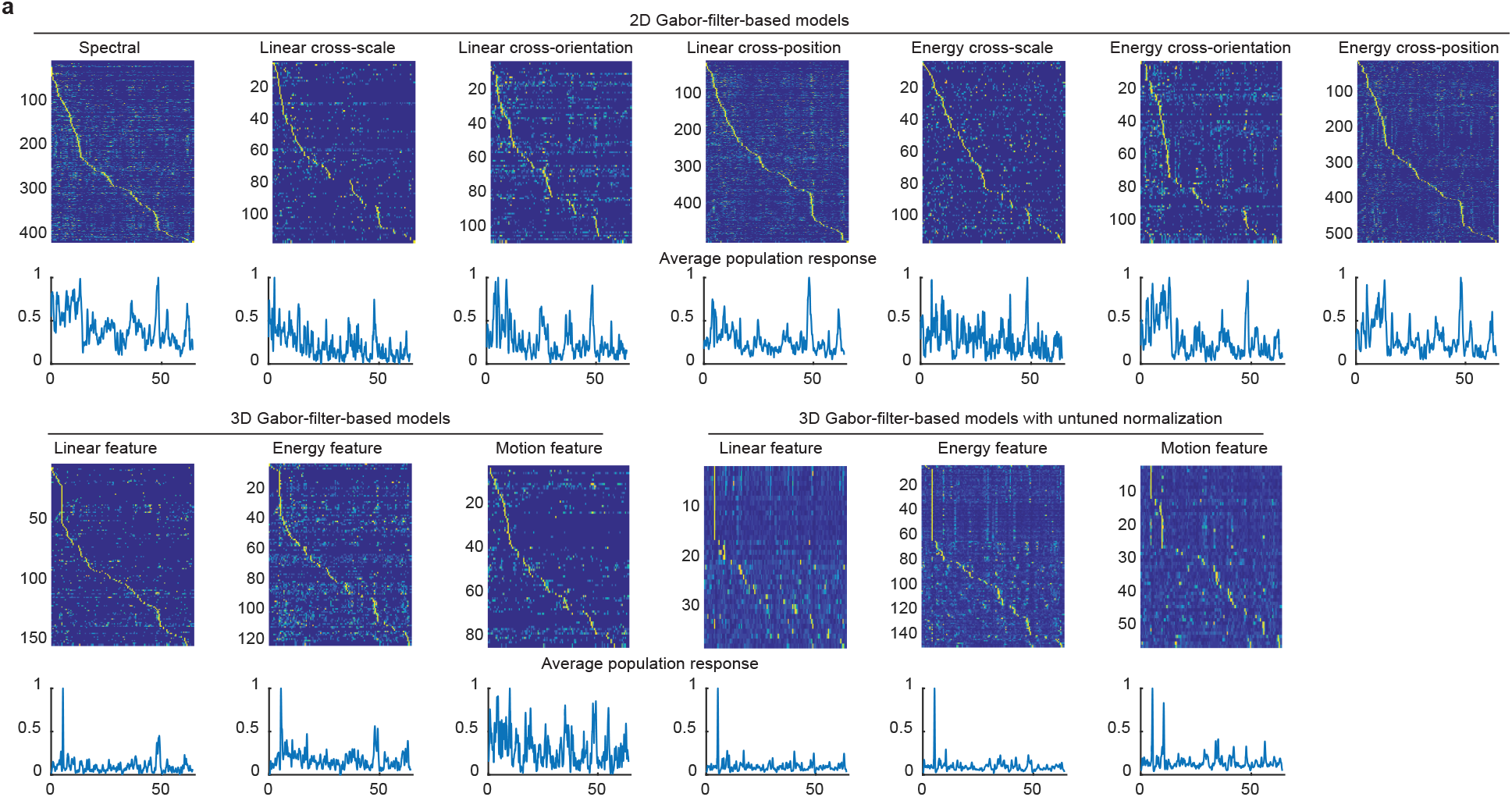
Encoding visual features of the naturalistic video using model neuron responses. (**a**) Simulated neuronal responses (PSTH) to the naturalistic video. The neurons were sorted by the timing of the strongest response. The color hue indicates the normalized value. The normalized average population response of each model was shown at the bottom.

**Supplementary Figure 12.**
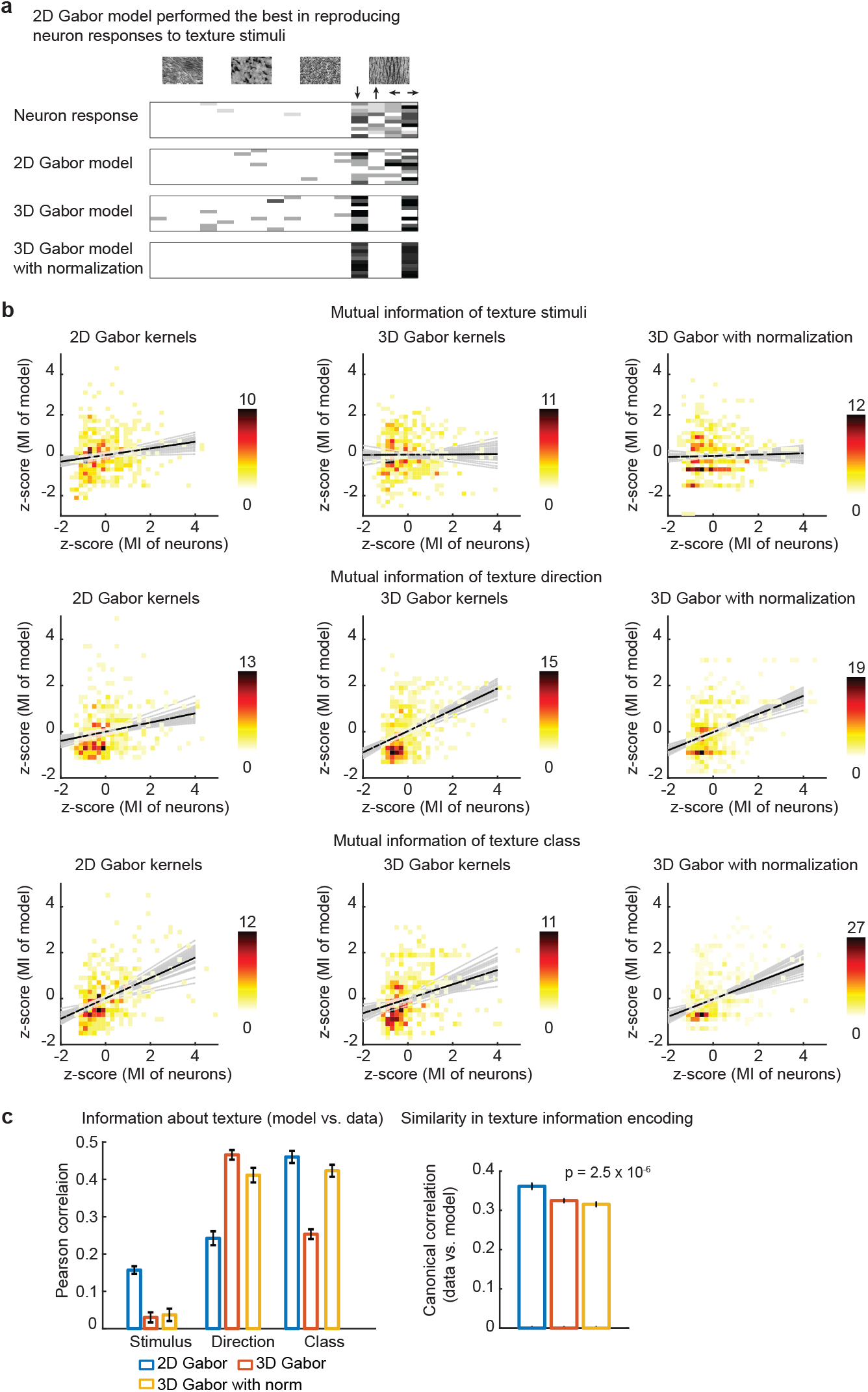
2D Gabor model performed the best in reproducing neuron responses to texture stimuli. (**a**) Model neuronal responses to texture stimuli. Binned spike count of a neuron, and its best 2G Gabor model, 3D Gabor model and 3D Gabor model with normalization. (**b**) Information about texture stimuli (top), texture direction (middle) and texture family (bottom) carried by neuron model vs. recorded neuronal responses. (**c**) Left, information about visual stimuli carried by neuron vs. model. Right, the similarity, measured by canonical correlation, of information encoding of texture stimuli between neuron models and neurons. Reported p-values are from a one-way ANOVA of mean canonical correlation values over all dimensions.

**Supplementary Figure 13.**
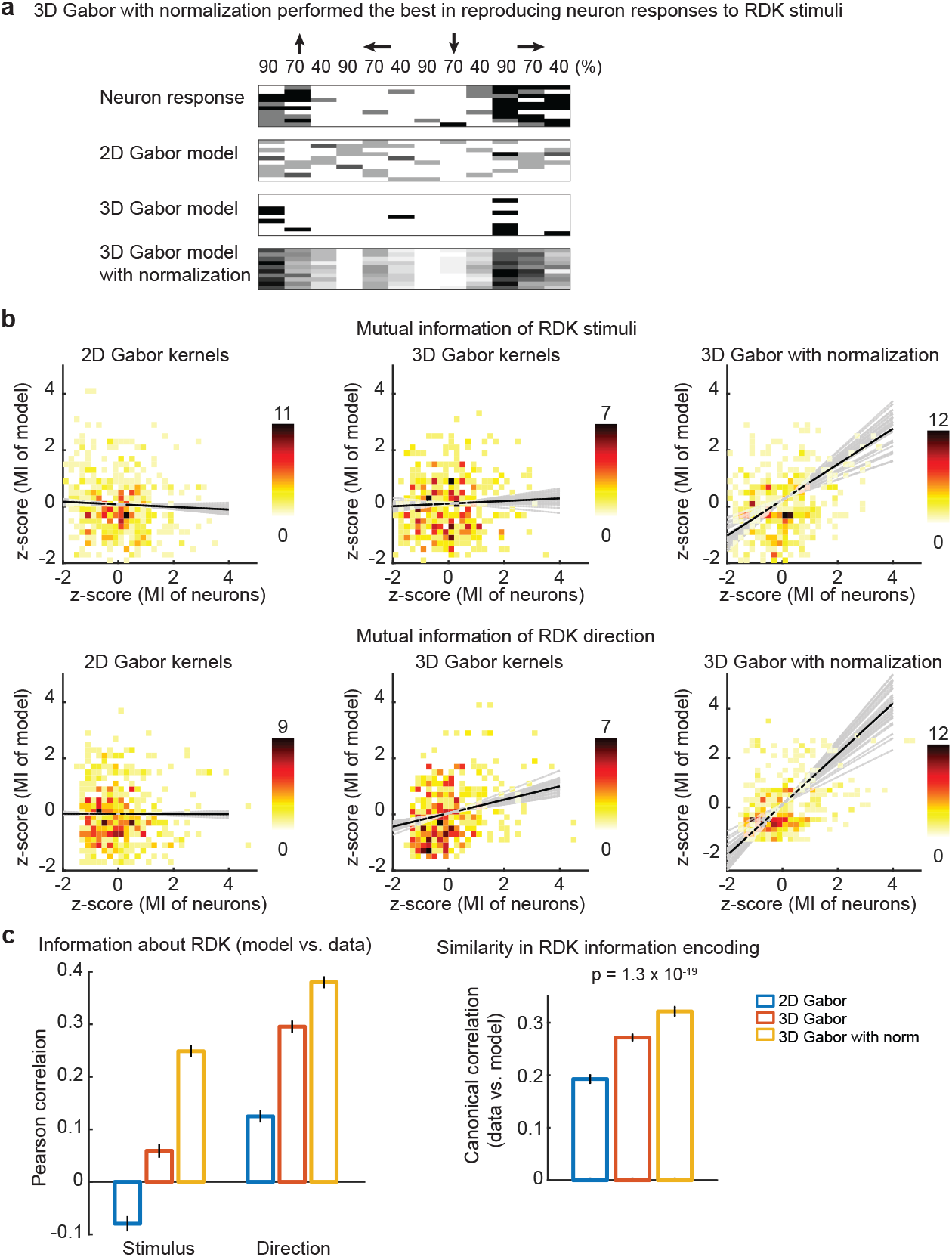
2D Gabor model performed the best in reproducing neuron responses to RDK stimuli. (**a**) Model neuronal responses to RDK stimuli. Binned spike count of a neuron, and its best 2G Gabor model, 3D Gabor model and 3D Gabor model with normalization. (**b**) Information about RDK stimuli (top), and RDK direction (bottom) carried by neuron model vs. recorded neuronal responses. (**c**) Left, information about visual stimuli carried by neuron vs. model. Right, the similarity, measured by canonical correlation, of information encoding of RDK stimuli between neuron models and neurons. Reported p-values are from a one-way ANOVA of mean canonical correlation values over all dimensions.

**Supplementary Figure 14.**
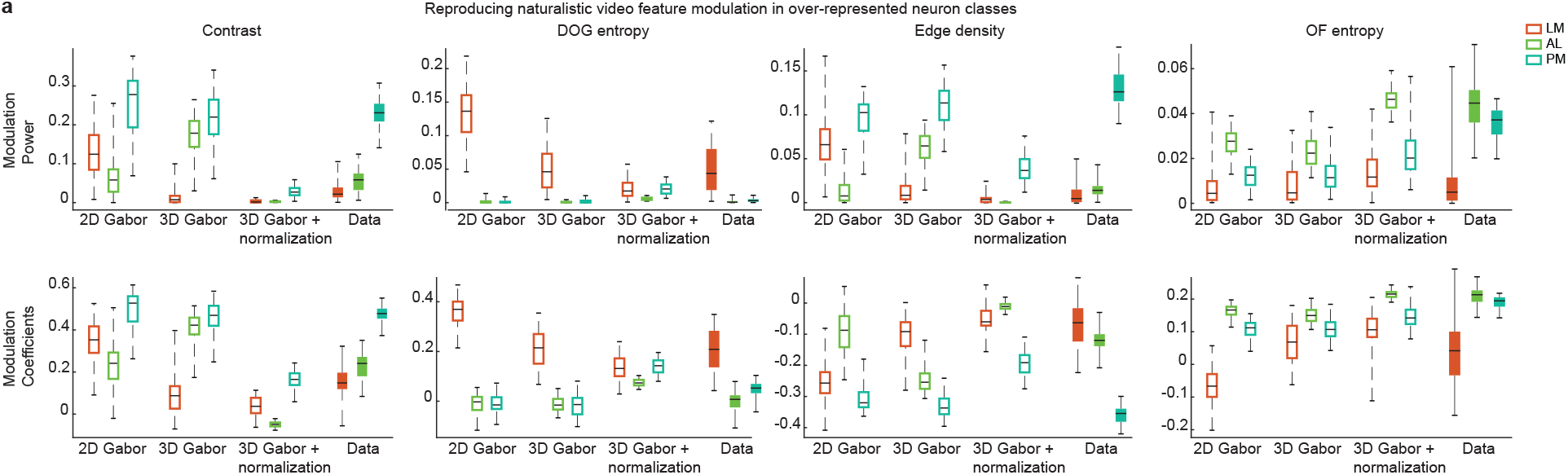
Reproducing the modulation of neuronal responses by visual features of naturalistic videos. (**a**) Boxplot of feature modulation power (upper) and modulation coefficients (down) of Gabor models for over-represented classes in HVAs (also, **Supplementary Fig. 7**).

## References

1. Park, J., Lee, G. & Chung, J. The PINK1-Parkin pathway is involved in the regulation of mitochondrial remodeling process. Biochem. Biophys. Res. Commun. 378, 518–23 (2009).

2. Tafazoli, S. et al. Emergence of transformation-tolerant representations of visual objects in rat lateral extrastriate cortex. Elife 6, 1–39 (2017).

3. Chang, L. & Tsao, D. Y. The Code for Facial Identity in the Primate Brain. Cell 169, 1013–1028.e14 (2017).

4. Baloh, R. H., Schmidt, R. E., Pestronk, A. & Milbrandt, J. Altered axonal mitochondrial transport in the pathogenesis of Charcot-Marie-Tooth disease from mitofusin 2 mutations. J. Neurosci. 27, 422–30 (2007).

5. Wang, Q., Sporns, O. & Burkhalter, A. Network Analysis of Corticocortical Connections Reveals Ventral and Dorsal Processing Streams in Mouse Visual Cortex. J. Neurosci. 32, 4386–4399 (2012).

6. Wang, Q., Gao, E. & Burkhalter, A. Gateways of Ventral and Dorsal Streams in Mouse Visual Cortex. J. Neurosci. 31, 1905–1918 (2011).

7. Berezovskii, V. K., Nassi, J. J. & Born, R. T. Segregation of feedforward and feedback projections in mouse visual cortex. J. Comp. Neurol. 519, 3672–3683 (2011).

8. Wang, Q. & Burkhalter, A. Area Map of Mouse Visual Cortex. J. Comp. Neurol. 502, 339– 357 (2007).

9. Siegle, J. H. et al. Survey of spiking in the mouse visual system reveals functional hierarchy. Nature 11–16 (2021).

10. Glickfeld, L. L., Andermann, M. L., Bonin, V. & Reid, R. C. Cortico-cortical projections in mouse visual cortex are functionally target specific. Nat. Neurosci. 16, 219–26 (2013).

11. Garrett, M. E., Nauhaus, I., Marshel, J. H. & Callaway, E. M. Topography and Areal Organization of Mouse Visual Cortex. J. Neurosci. 34, 12587–12600 (2014).

12. Kalatsky, V. A., Stryker, M. P. & Foundation, W. M. K. New paradigm for optical imaging: Temporally encoded maps of intrinsic signal. Neuron 38, 529–545 (2003).

13. Schuett, S., Bonhoeffer, T. & Hübener, M. Mapping retinotopic structure in mouse visual cortex with optical imaging. J. Neurosci. 22, 6549–6559 (2002).

14. Hubel, D. H. & Wiesel, T. AND FUNCTIONAL ARCHITECTURE IN THE CAT ’ S VISUAL CORTEX From the Neurophysiolojy Laboratory, Department of Pharmacology central nervous system is the great diversity of its cell types and inter-receptive fields of a more complex type ( Part I ) and to. J. Physiol. 160, 106–154 (1962).

15. Jin, M. & Glickfeld, L. L. Mouse Higher Visual Areas Provide Both Distributed and Specialized Contributions to Visually Guided Behaviors. Curr. Biol. 30, 4682–4692.e7 (2020).

16. Stringer, C., Michaelos, M., Tsyboulski, D., Lindo, S. E. & Pachitariu, M. High-precision coding in visual cortex. Cell 184, 2767–2778.e15 (2021).

17. Andermann, M. L., Kerlin, A. M., Roumis, D. K., Glickfeld, L. L. & Reid, R. C. Functional Specialization of Mouse Higher Visual Cortical Areas. Neuron 72, 1025–1039 (2011).

18. Marshel, J. H., Garrett, M. E., Nauhaus, I. & Callaway, E. M. Functional specialization of seven mouse visual cortical areas. Neuron 72, 1040–1054 (2011).

19. Goltstein, P. M., Reinert, S., Bonhoeffer, T. & Hübener, M. Mouse visual cortex areas represent perceptual and semantic features of learned visual categories. Nat. Neurosci. 24, (2021).

20. Han, X., Vermaercke, B. & Bonin, V. Cellular organization of visual information processing channels in the mouse visual cortex. BioRxiv (2020).

21. Juavinett, A. L. L., Callaway, E. M. M., Information, S., Juavinett, A. L. L. & Callaway, E. M. M. Pattern and Component Motion Responses in Mouse Visual Cortical Areas. Curr. Biol. 25, 1759–1764 (2015).

22. Bolaños, F., Orlandi, J. G., Jagadeesh, A. V, Gardner, J. L. & Benucci, A. Processing of visual textures in the primary and secondary visual cortex of the mouse Non-human primate work. in Society for Neuroscience Annual Meeting 1–12 (2021).

23. Sit, K. K. & Goard, M. J. Distributed and retinotopically asymmetric processing of coherent motion in mouse visual cortex. Nat. Commun. 11, 1–14 (2020).

24. Freeman, J., Ziemba, C. M., Heeger, D. J., Simoncelli, E. P. & Movshon, J. A. A functional and perceptual signature of the second visual area in primates. Nat. Neurosci. 16, 974–981 (2013).

25. DiCarlo, J. J., Zoccolan, D. & Rust, N. C. How does the brain solve visual object recognition? Neuron 73, 415–434 (2012).

26. Ziemba, C. M., Freeman, J., Movshon, J. A. & Simoncelli, E. P. Selectivity and tolerance for visual texture in macaque V2. Proc. Natl. Acad. Sci. 113, E3140–3149 (2016).

27. Khawaja, F. A., Liu, L. D. & Pack, C. C. Responses of MST neurons to plaid stimuli. J. Neurophysiol. 110, 63–74 (2013).

28. Jeffrey N. Stirman, Ikuko T. Smith, Michael W. Kudenov, S. L. S. Wide field-of-view, multi-region two-photon imaging of neuronal activity. Nat. Biotechnol. 34, 857–862 (2016).

29. Madisen, L. et al. Transgenic Mice for Intersectional Targeting of Neural Sensors and Effectors with High Specificity and Performance. Neuron 85, 942–958 (2015).

30. Chen, T. W. et al. Ultrasensitive fluorescent proteins for imaging neuronal activity. Nature 499, 295–300 (2013).

31. Smith, I. T., Townsend, L. B., Huh, R., Zhu, H. & Smith, S. L. Stream-dependent development of higher visual cortical areas. Nat. Neurosci. 20, 200–208 (2017).

32. Prusky, G. T. & Douglas, R. M. Characterization of mouse cortical spatial vision. Vision Res. 44, 3411–3418 (2004).

33. Duan, H. & Wang, X. Visual attention model based on statistical properties of neuron responses. Sci. Rep. 5, 8873 (2015).

34. Goldbach, H. C., Akitake, B., Leedy, C. E. & Histed, M. H. Performance in even a simple perceptual task depends on mouse secondary visual areas. Elife 10, 1–39 (2021).

35. Carandini, M., Heeger, D. J. & Movshon, J. A. Linearity and Normalization in Simple Cells of the Macaque Primary Visual Cortex. J. Neurosci. 17, 8621–8644 (1997).

36. Yoshida, T. & Ohki, K. Natural images are reliably represented by sparse and variable populations of neurons in visual cortex. Nat. Commun. 11, (2020).

37. Okazawa, G., Tajima, S. & Komatsu, H. Image statistics underlying natural texture selectivity of neurons in macaque V4. Proc. Natl. Acad. Sci. 112, E351–360 (2014).

38. Cowley, B. R., Smith, M. A., Kohn, A. & Yu, B. M. Stimulus-Driven Population Activity Patterns in Macaque Primary Visual Cortex. PLoS Comput. Biol. 12, 1–31 (2016).

39. de Vries, S. E. J. et al. A large-scale standardized physiological survey reveals functional organization of the mouse visual cortex. Nat. Neurosci. 23, 138–151 (2020).

40. Britten, K. H., Newsome, W. T., Shadlen, M. N., Celebrini, S. & Movshon, J. A. A relationship between behavioral choice and the visual responses of neurons in macaque MT. Vis. Neurosci. 13, 87–100 (1996).

41. Saleem, A. B. Two stream hypothesis of visual processing for navigation in mouse. Curr. Opin. Neurobiol. 64, 70–78 (2020).

42. Smith, S. L. & Häusser, M. Parallel processing of visual space by neighboring neurons in mouse visual cortex. Nat. Neurosci. 13, 1144–1149 (2010).

43. Pachitariu, M., et al. Suite2p: beyond 10,000 neurons with standard two-photon microscopy. bioRxiv 061507 (2017).

44. Harris, K. D., Quiroga, R. Q., Freeman, J. & Smith, S. L. Improving data quality in neuronal population recordings. Nat. Neurosci. 19, 1165–1174 (2016).

45. Pnevmatikakis, E. A., Merel, J., Pakman, A. & Paninski, L. Bayesian spike inference from calcium imaging data. Adv. Neural Inf. Process. Syst. 26, 1250–1258 (2013).

46. Portilla, J. & Simoncelli, E. P. Parametric texture model based on joint statistics of complex wavelet coefficients. Int. J. Comput. Vis. 40, 49–71 (2000).

47. Stirman, J. N., Townsend, L. B. & Smith, S. L. A touchscreen based global motion perception task for mice. Vision Res. 127, 74–83 (2016).

48. Canny, J. A Computational Approach to Edge Detection. IEEE Trans. Pattern Anal. Mach. Intell. PAMI-8, 679–698 (1986).

49. Hermundstad, A. M. et al. Variance predicts salience in central sensory processing. Elife 3, 1–28 (2014).

50. Yu, Y., Burton, S. D., Tripathy, S. J. & Urban, N. N. Postnatal development attunes olfactory bulb mitral cells to highfrequency signaling. J. Neurophysiol. 114, 2830–2842 (2015).

51. Froudarakis, E. et al. Population code in mouse V1 facilitates readout of natural scenes through increased sparseness. Nat. Neurosci. 17, 851–857 (2014).

52. Baden, T. et al. The functional diversity of retinal ganglion cells in the mouse. Nature 529, 345–350 (2016).

53. Adelson, E. H. & Bergen, J. R. Spatiotemporal energy models for the perception of motion. J. Opt. Soc. Am. A 2, 284 (1985).

